# *TMPRSS2-ERG* confers resistance to antiandrogens: mechanism and therapeutic implications

**DOI:** 10.1101/2025.01.17.633582

**Authors:** Arunachalam Sekar, Rishita Chatterjee, Boobash Raj Selvadurai, Lipika R. Pal, Aakanksha Verma, Nishanth Belugali Nataraj, Diana Drago Garcia, Suvendu Giri, Alessandro Genna, Feride Karatekin, Nitin Gupta, Mirie Zerbib, Yaron Vinik, Tamir Avioz, Eviatar Weizman, Eyal David, Alejändro A. Schaffer, Yunqian Pan, Haojie Huang, Wytske M. van Weerden, Eva Corey, Hazel Hunt, Andrew E. Greenstein, Ronnie Blecher-Gonen, Roni Oren, Ariel Afek, Ido Amit, Sima Lev, Anson Ku, Sumeyra Kartal, Jack R. Bright, Rosina T. Lis, William L. Dahut, Adam G. Sowalsky, Eytan Ruppin, Yosef Yarden

**Author notes:** To whom correspondence should be addressed: Yosef Yarden, Department of Immunology and Regenerative Biology, Weizmann Institute of Science, Rehovot 76100, Israel. **Current addresses:** Bugworks Research Inc, C-CAMP, NCBS campus, Bangalore-560065, India. The Francis Crick Institute, London NW1 1AT, United Kingdom (Diana Drago Garcia). National University of Singapore, Institute of Molecular and Cell Biology (IMCB), Singapore 138673. **Lead contact:** Yosef Yarden.

## Abstract

Approximately 50% of prostate cancer (PCa) patients harbor fusions involving the *TMPRSS2* and *ERG* genes. Despite this, tailored therapies targeting the fused gene, *tERG*, remain undeveloped. Our study analyzed biopsy samples from two clinical trials assessing the efficacies of androgen receptor (AR) signaling inhibitors (ARSIs). The results revealed that *tERG* promotes resistance to ARSIs and is associated with elevated levels of the glucocorticoid receptor (GR). Subsequent assays showed that GR directly interacts with tERG, alleviates allosteric autoinhibition and prevents chemotherapy-induced tERG degradation. In PCa models, either inhibiting GR or lowering cortisol levels suppressed tumor growth in tERG-positive models, but not in fusion-negative models. In addition, patient-derived fusion-positive xenografts displayed enhanced sensitivity to combined GR and AR inhibitors. Collectively, these findings highlight *TMPRSS2-ERG* as a new biomarker and propose that simultaneous inhibition of GR and AR may specifically benefit *tERG*-positine patients. However, GR stimulatory corticosteroid therapies may not be advisable for this patient subgroup.

## Introduction

Prostate cancer (PCa) is the second most common cause of cancer-related deaths in men. The majority of tumors fall into one of seven subtypes defined by specific gene fusions or mutations ^1^. More than 50% of Caucasian patients with PCa harbor chromosomal rearrangements in genes encoding for specific ETS (E26 transformation-specific) family transcription factors (TFs) ^2–5^. The most common rearrangement fuses the promoter region of the androgen-regulated gene *TMPRSS2* (transmembrane protease, serine 2), with the coding region of *ERG*, a member of the ETS family, which lies near *TMPRSS2* on human chromosome 21. Similar but less frequent ETS fusions engage *ETV1*, *ETV4* or *ETV5,* which are on other chromosomes. One outcome of the *TMPRSS2-ERG (tERG)* gene fusion is androgen-inducible overexpression of the ERG protein. Notably, in normal benign prostatic tissue and benign prostatic hyperplasia *TMPRSS2* fusion genes do not form. However, *TMPRSS2-ERG* gene fusions are observed at the earliest stages of PCa development, including a subset of high-grade prostatic intraepithelial neoplasia^6^. When present, the fusion gene is highly expressed in both primary and castration-resistant tumors ^4,7^. Although *tERG* is associated with high Gleason scores and the presence of *TMPRSS2-ERG* fusion marks relatively aggressive tumors, the role for *tERG* as a prognostic marker is complex.

ERG and other members of the ETS family share a well-conserved DNA binding element, ETS, which binds with the DNA motif GGA(A/T) ^8,9^. In addition to the ETS domain, a subset of ETS proteins harbor the pointed domain (PNT), which facilitates protein-protein interactions and dimerization. tERG drives unique transcriptional programs, represses prostate epithelial differentiation genes, such as *PSA* and *SLC45A3*, and controls several downstream targets, including components of the NOTCH signaling pathway, MYC, EZH2 and WNT ^10–13^. Through binding with coactivators and corepressors, the transcriptional activity of ETS proteins is stringently modulated ^14^. For example, tERG can physically interact with BAF (SWI/SNF) complexes ^15^, to regulate several cellular processes in PCa cells, including gene expression and ERG-driven basal-to-luminal transition. In addition, the *tERG* fusion gives rise to various N-terminally truncated ERG proteins ^16^. The majority of the truncated fusions are resistant to degradation because they lack the degron motif for a ubiquitin ligase encoded by the *SPOP* gene, which is mutated in approximately 11% of PCa ^17,18^. There is an analogous ERG-inhibitory mechanism carried out by the ETS2 repressor factor, ERF, which undergoes inactivation in PCa by means of recurrent point mutations and focal deletions ^19^. Thus, hormonal control of ERG expression, competition with ERF and avoidance of degradation offer therapeutic opportunities for targeting ERG in PCa.

The currently approved therapies for castration-resistant prostate cancer (CRPC) include both non-steroidal anti-androgens (NSAA, e.g., enzalutamide) and inhibitors of CYP17A1 (e.g., abiraterone), a cytochrome P450 enzyme that catalyzes production of several steroid hormones, including androgens. Unfortunately, the initial positive responses to these therapies are typically followed by drug resistance and tumor relapses. Several studies have shown that activation of the glucocorticoid receptor (GR) permits resistance to enzalutamide ^20–24^. Both the endogenous steroid ligand of GR, cortisol, and synthetic GR ligands, such as dexamethasone (DEX), transduce their actions by binding to GR. GR activity is able to replace AR due to similar promoter sequences and downstream targets, including the anti-apoptotic mediators *DUSP1* and *SGK1.* Normally, GR expression is silenced by AR ^23^. Hence, prolonged inhibition of AR can result in GR upregulation, underscoring the selective pressure to maintain downstream target gene expression.

Earlier, we demonstrated that FLI1, along with other ETS family members, directly interacts with GR ^25^. Furthermore, inhibiting GR in Ewing sarcoma models—pediatric tumors driven by the *EWS-FLI1* fusion gene—led to decreased tumor growth and metastasis. Given these findings and GR’s role in resistance to AR signaling inhibitors (ARSIs) ^20^, we explored ETS-driven PCa. Data from two clinical trials revealed that PCa patients expressing *tERG*-positive fusions frequently exhibit resistance to antiandrogens, and this is linked to elevated GR levels after treatment. We also discovered that GR binds to ERG’s DNA-binding domain, stabilizing ERG and mitigating an intrinsic mechanism of allosteric autoinhibition. As a result, ERG’s transcriptional activity increases when it forms complexes with GR. In patient-derived xenografts, *tERG*-positive models demonstrated heightened sensitivity to GR inhibition compared to *tERG*-negative models. These findings identify fused *ETS* genes as a novel class of PCa biomarkers, predicting resistance to second-generation antiandrogens, and suggesting that corticosteroid therapy may be contraindicated for patients with PCa expressing an *ETS* fusion gene.

## Results

### Tumoral tERG predicts resistance of patients with prostate cancer to anti-androgen treatments

We began by re-analyzing data collected by a recent PCa clinical trial, National Cancer Institute (NCI) 15-c-0124 (NCT02430480). This study performed immunohistochemical (IHC) analysis, whole-exome and RNA-sequencing on biopsy samples from 37 men with prostate cancer, before they received ADT plus enzalutamide for 6 months ^26,27^. As expected, there is bimodal distribution of ERG’s mRNA abundance in the pre-treatment data, and this corresponded to the presence of *ERG* fusions in the high IHC group (Figs. 1A and S1A). Due to upregulation by androgens, ERG levels were significantly altered post antiandrogen treatment: reduced in the positive tERG patients (p=5.19E-07, one sided Wilcoxon test) and increased in the negative tERG patients (p=0.00037; Fig. 1B). Notably, along with AR and ERG, GR (encoded by the *NR3C1* gene) has been reported to play an important role in PCa ^20,28,29^. In contrast to ERG, GR displayed a unimodal pre-treatment distribution but, irrespective of ERG’s status, GR’s IHC increased substantially post treatment (Figs. 1A and 1B; lower panels), likely due to relief of the previously reported AR-mediated suppression of GR ^21,23^. Quite surprisingly, the majority of patients with tERG and high *ERG* expression at baseline exhibited resistance to the antiandrogen treatment (p=0.0047; one-sided Wilcoxon test; Fig. S1B). Treatment resistance was also associated with high post-treatment ERG, as determined using IHC (p=0.0073; Fig. 1C; upper panel), as well as with high post-treatment GR levels, which were determined using either mRNA abundance (p=0.039; Fig. 1C, lower panel) or IHC (p=0.018; Fig. S1C). Together, these observations uncovered functional association between the mutant form of ERG, high GR and resistance to enzalutamide.

**Figure 1:**
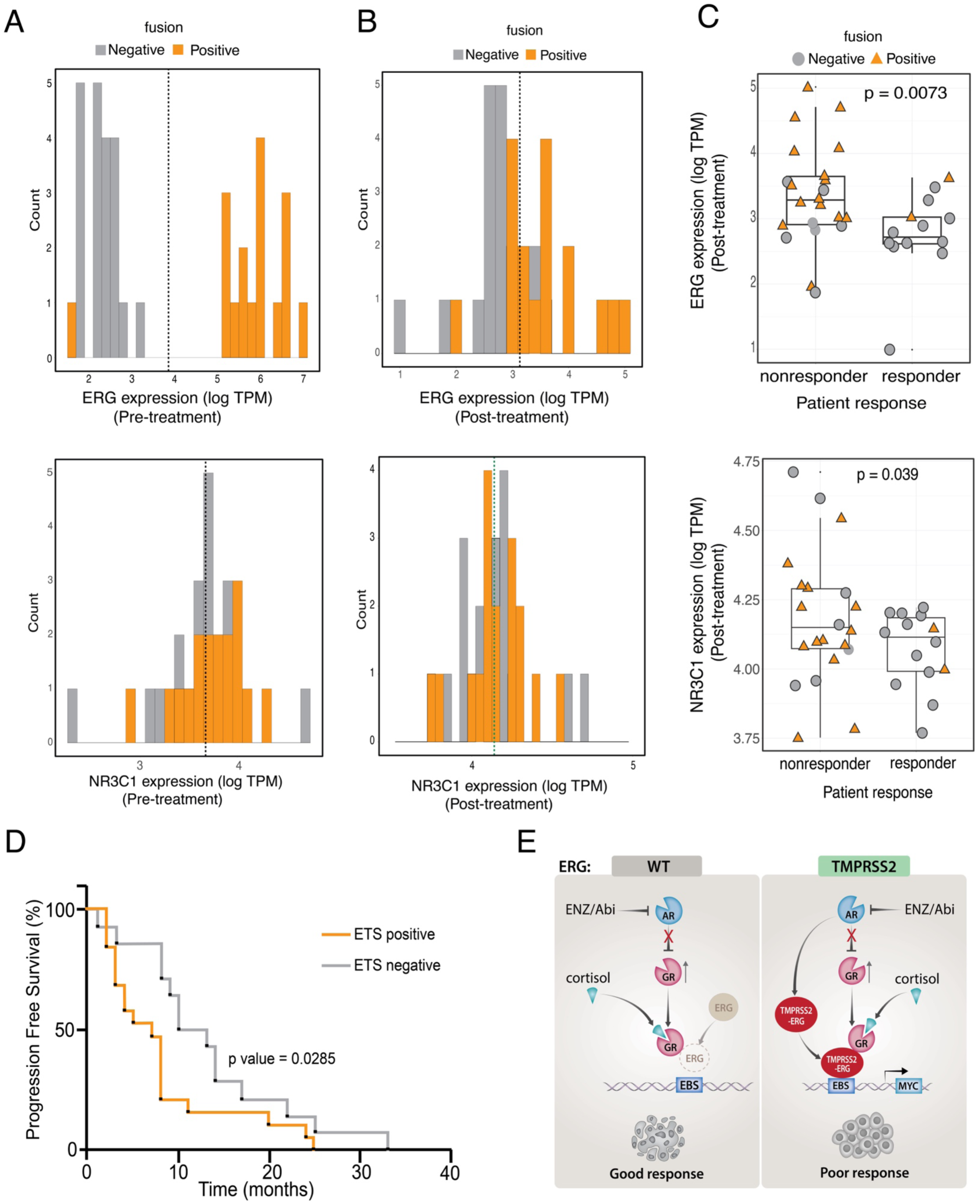
The status of ERG (wildtype or rearranged) and GR abundance predict the responses of patients with prostate cancer to ADT plus enzalutamide (see also Figure S1). (**A** and **B**) Biopsies were obtained from 37 men with intermediate-to high-risk prostate cancer before receiving ADT plus enzalutamide for 6 months. The tissues were used for IHC, whole exome sequencing and RNA-sequencing. Shown are the distributions of pre-(A) and post-treatment (B) levels of ERG (upper panels) and GR (NR3C1; lower panels) in all patients. Fusion status of ERG in patients is shown in orange for positive fusion (tERG) and in grey for negative fusion (wildtype ERG). (**C**) Post-treatment ERG and GR expression levels are presented against patient response to treatment with ADT plus enzalutamide. The P-values presented were determined using either one-sided Wilcoxon (ERG) or one-sided unpaired T-test (GR). (**D**) Data were derived from the control arm of clinical trial NCT01576172 (NCI 9012). All patients included in this arm (n=58) were treated with abiraterone plus prednisone. Biopsy samples were obtained from metastatic sites and analyzed using RNA-seq and immunohistochemistry (for GR and ERG). Shown are progression free survival (PFS) curves corresponding to the tERG negative and tERG positive groups (p=0.0285; Gehan-Breslow-Wilcoxon two sided test). (**E**) A working model depicting the inferred functional interaction between TMPRSS2-ERG and the hormone-activated glucocorticoid receptor (GR) in prostate cancer. Normallly, AR suppresses GR in tumors driven by the androgen receptor (AR). However, by inhibiting AR signaling, drugs like enzalutamide (ENZ) and abiraterone (Abi) relieve this repression, hence they up-regulate GR. In approximately 50% of prostate tumors, the androgen-inducible *TMPRSS2* gene is fused to ERG. The fused gene, *TMPRSS2-ERG* (*tERG*), regulated by AR and the encoded protein might form stable complexes with active GRs. However, this may not occur in fusion-negative tumors. Once formed, tERG-GR complexes bind with the ERG binding site (EBS) on DNA to stimulate transcription of *MYC* and other oncogenic targets. This mechanism can explain why *TMPRSS2-ERG*-positive tumors might gain resistance to anti-androgen treatments.

Next, we asked if the correlation between tERG expression and lack of response to anti-androgen treatment extends to metastatic castration-resistant prostate cancer (mCRPC). For this, we analyzed the results of another clinical trial, NCT01576172 (NCI 9012) ^30^. All patients of this trial underwent metastatic site biopsy, stratified by the status of ERG and ETV1 (using IHC and in situ hybridization, respectively) and randomly assigned to abiraterone plus prednisone, either without (arm A) or with an inhibitor of poly(ADP-ribose) polymerase-1 (PARP1), veliparib (arm B). We focused on a subset of arm A patients (N=58) for whom survival data was available. These patients presented samples with matching RNA-seq, GR and ERG immunohistochemistry, and their PSA responses were available. A summary Kaplan-Meier survival analysis is presented in Figure 1D. As predicted, the median progression free survival (PFS) of the *tERG*-negative group (11.5 months) was significantly longer (p=0.0285; Gehan-Breslow-Wilcoxon test) than the PFS rate of the *tERG*-positive cohort (7.0 months).

In conclusion, according to the analysis of the results reported by two independent clinical trials (total, 95 patients), expression of the fused form of the *ERG* gene associates with poorer response of patients with PCa to treatments with two different ARSIs, enzalutamide and abiraterone plus prednisone.

### The association of *tERG* with resistance to ARSIs and relatively high GR offers a working model

To further examine potential clinical associations between the levels of ERG and GR, we analyzed two post-treatment datasets: a microarray set, GSE102124 ^31^, derived from 19 patients previously treated with ADT plus abiraterone (Fig. S1D), and an RNA-seq dataset ^32,33^, derived from 160 patients that were treated with front-line ADT (Fig. S1E). The presented scatter plots and respective P-values, 0.016 and 0.00039, further supported the existence of positive association between ERG and GR in clinical specimens. Taken together with the previosly reported stimulatory interactions between GR and another ETS family member, FLI1 ^25^, the data shown in Figures 1 and S1 suggest the following working model (see Figure 1E): Under normal conditions, GR expression is suppressed by androgen signaling ^21,23,34^. However, ARSIs like enzalutamide (ENZ) and abiraterone (Abi) can lift this suppression and increase GR expression. In tumors positive for the *TMPRSS2-ERG* fusion, GR may directly interact with tERG, in similarity to the reported GR-FLI1 complexes. Once the putative GR-tERG complexes are formed, they bind to the ERG binding site (EBS) on DNA, thereby promoting the transcription of oncogenic targets like MYC. Notably, upregulation of GR can bypass AR signaling by activating a partly overlapping set of target genes ^20,35^. According to our results, this predicted sequence of events may not occur in *TMPRSS2*-negative tumors, which typically well respond to anti-androgen treatments (Fig. 1C).

### GR forms druggable complexes with oncogenic ETS proteins

In anticipation of direct interactions between ERG and the hormone-bound form of GR, we employed the previosly described protein complementation assay (reviewed in ^36^). The modified protocol we used utilized inactive fragments of Gaussia luciferase (Gluc), which was split into an amino-terminal fragment, Gluc1, and a carboxyl-terminal fragment, Gluc2. Firstly, we constructed a library of Gluc1 fusion proteins that included the ETS proteins frequently rearranged in PCa (i.e., ERG, ETV1, ETV4 and ETV5), along with ETV6, as a negative control. The levels of expression of the Gluc proteins are shown in Figure S2A. Note that Gluc1-ETV5 displayed relatively low expression. The normalized signals presented in Figure 2A indicated that stimulation with DEX can significantly increase the interaction between GR and all ETS proteins we analyzed, except for ETV6. In addition, the hormone-induced interactions were inhibited when cells were treated with a combination of DEX and RU486 (mifepristone), a GR antagonist.

**Figure 2:**
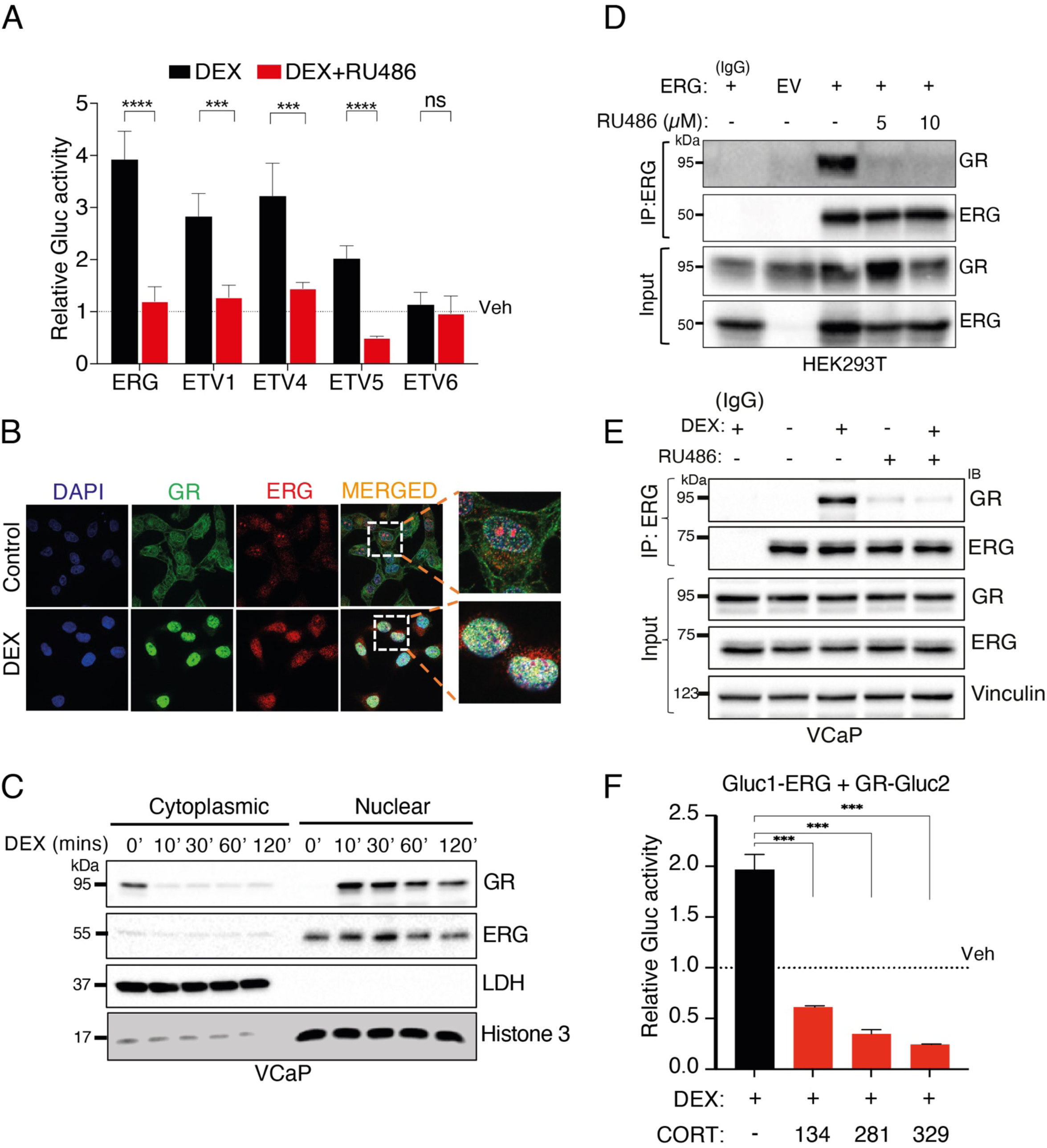
The physical interactions between GR and ERG take place in the nucleus and they can be disrupted by a GR antagonist (see also Figure S2). (**A**) HEK293T cells growing in charcoal-stripped serum (CSS) were transfected (in sextuplicates) with GR-Gluc2, along with the indicated Gluc1-ETS plasmids. Twenty-four hours post transfection, cells were treated with vehicle, DEX (1 μM), or a combination of DEX and RU486 (1 μM). Shown are normalized fold changes in luminescence. ***, p < 0.001; ****, p < 0.0001; ns, not significant. Note that ETV6 was used as a negative control. (**B**) CSS grown HeLa cells were stimulated with either DEX or vehicle (control) for 10 minutes. Thereafter, cells were fixed and processed for immunofluorescence analysis that used DAPI (blue; DNA staining), and antibodies specific to GR (green) or ERG (red). The framed areas in the merged column are magnified in the rightmost column. Bar, 20 μm. (**C**) Serum-starved VCaP cells expressing tERG were treated with DEX (1 μM) for the indicated time intervals. Untreated and DEX-treated cells were subjected to subcellular fractionation and immunoblotting for GR and ERG. LDH and histone 3 were used as cytoplasmic and nuclear markers, respectively. (**D**) HEK293T cells were transfected with an empty vector (EV) or with a plasmid encoding ERG, and 24 hours post transfection cells were treated with RU486 at the indicated concentrations, for 8 hours. Thereafter, we subjected cell extracts to immunoprecipitation with an anti-ERG antibody or with a control antibody (IgG), and immunoblotted with the indicated antibodies. A fraction (5%) of the lysate was used as input control. (**E**) Serum-starved VCaP cells were treated for 60 minutes with vehicle, DEX (1 μM), RU486 (1 μM), or the combination of drugs. Extracts were processed for immunoprecipitation (IP) and immunoblotting (IB). Note that two different anti-ERG antibodies were used, one from Cell Signalling Technology (CST) and the other from Santa Cruz Biotechnology (SCB). IgG, control antibody. (**F**) Serum-starved VCaP cells were treated for 60 minutes with DEX (1 μM) in the absence or presence of the following selective GR modulators (SGRMs, 10 μM): CORT125134 (C134, relacorilant), CORT125281 (C281, exicorilant) and CORT125329 (C329). Luminescence was determined in biological triplicates. All experiments were repeated thrice. ***, p<0.001.

Next, we visualized the interaction between GR and ERG at the subcellular level. Due to low signal to noise ratio observed with PCa cells, we utilized HeLa cells that were pre-starved, stimulated with DEX and then examined. Once stimulated with the steroid hormone, the endogenous GR rapidly translocated to the nucleus (Fig. 2B). In parallel, the diffusely distributed nuclear ERG redistributed into condensed intra-nuclear structures that contained the translocated GR. However, some ERG-containing puncta, which were devoid of GR, acquired polarized peripheral locations of an unknown nature. Subcellular fractionation corroborated these observations (Fig. 2C): as early as 10 minutes after exposure to DEX, the initially cytoplasmic GRs translocated to the nucleus, which already contained most of the cellular ERG molecules. Notice that we used a nuclear marker along with a cytoplasm-specific marker and observed early DEX-dependent enhancement of the GR protein and late downregulation of this receptor, due to the previously described ‘homologous downregulation’ ^37^.

To directly validate the physical interactions between GR and tERG, we carried out co-immunoprecipitation assays (Fig. 2D). In untreated HEK293T cells ectopically expressing ERG or an empty vector (EV), we observed high levels of the endogenous GR protein that was pulled-down with an anti-ERG antibody. Consistent with specific interactions, treatment with RU486 abolished complex formation. Similarly, in ERG immunoprecipitates obtained from tERG-expressing PCa cells (VCaP) ^38^, which were pre-starved for steroid hormones, DEX increased the amount of GR in ERG immunoprecipitates (Fig. 2E). Although *TMPRSS2*-*ETV4* fusions are less frequently overexpressed than *TMPRSS2-ERG* among human patients, increased abundance of ETV4 promotes tumor growth in xenograft models ^39,40^. As expected, analysis of PC3 cells, which naturally overexpress ETV4, confirmed physical GR-ETV4 associations that were increased by DEX and decreased by RU486 (Fig. S2B). Because non-steroidal GR antagonists hold promise for PCa treatment ^41^, we examined a series of three non-steroidal selective GR antagonists (from Corcept Therapeutics). In similarity to RU486, each of the three antagonists inhibited the ability of DEX to drive formation of the GR-ERG complex (Fig. S2C), as well as reduced the luminescence signals obtained when combined with DEX (Fig. 2F). In summary, following stimulation with DEX GR translocates to the nucleus and forms non-covalent bonds with ERG, but these interactions can be inhibited using pharmacological agents.

### The DNA binding domain of GR and the ETS domain of ERG are engaged in complex formation

Similar to other ETS family members, ERG harbors an ∼80-residue domain, PNT, and a conserved ETS domain, which binds DNA with preference for an invariant 5′-GGA(A/T)-3′ core ^42^. GR also contains a DNA binding domain (DBD), an N-terminal modulatory domain (NTD), a hinge region (HR) and a C-terminal ligand binding domain, LBD ^43^. To map the domains mediating the interaction between GR and ERG, we made use of several deletion mutants of each protein. The GR deletion mutants schematically presented in Figure S2D were cloned into the Gluc2 plasmid and expressed in both DU145 (Fig. S2E) and HEK293T cells (Fig. S2F), along with either full-length HA-ERG (Fig. S2E) or Gluc1-ERG (Fig. S2F). The results of the co-immunoprecipitation and protein complementation assays revealed that the region comprising DBD+HR+LBD (mutant denoted GR_3) generated slightly stronger interaction signals than the full-length GR (Figs. S2E and S2F). No protein complementation signal was observed when using another GR deletion mutant that included HR+LBD. Hence, we concluded that the DBD of GR mediates the interaction with ERG. Similar analysis of a set of ERG deletion mutants (Fig. S2G; see list of primers in Table S1) indicated that the ETS domain is critical for the interaction (Figs. S2H and S2I). In conclusion, the DNA-binding domain of GR forms a physical complex with ETS, the DNA binding domain of ERG.

### GR protects tERG from basal and chemotherapy-induced degradation

Ubiquitination and degradation of the ERG oncoprotein suppress PCa progression, while resistance to degradation serves as a mechanism that contributes to the elevation of truncated ERG proteins in PCa ^17,18,44^. Hence, we set out to investigate whether by forming a physical complex with GR, tERG gains stability. To test this model, we treated VCaP cells with RU486. As shown in Figure 3A and 3B, RU486 treatment reduced the abundance of the tERG protein in a dose-and time-dependent manner. Notably, neither DEX nor RU486 modified the abundance of mRNAs encoding tERG, but treatment with enzalutamide reduced tERG’s transcript levels, as expected (Fig. S3A). To further rule out transcription-mediated regulation of tERG by GR, we examined the fate of a constitutively overexpressed (androgen-independent) tERG. To this end, we transfected the ERG-low DU145 cells with a tERG-encoding plasmid and established two cell lines (Fig. S3B). Using one of the two stable derivatives we confirmed RU486-induced destabilization of tERG (Fig. S3C), consistent with the possibility that GR posttranslationally regulates tERG by means of protein-protein interactions.

**Figure 3:**
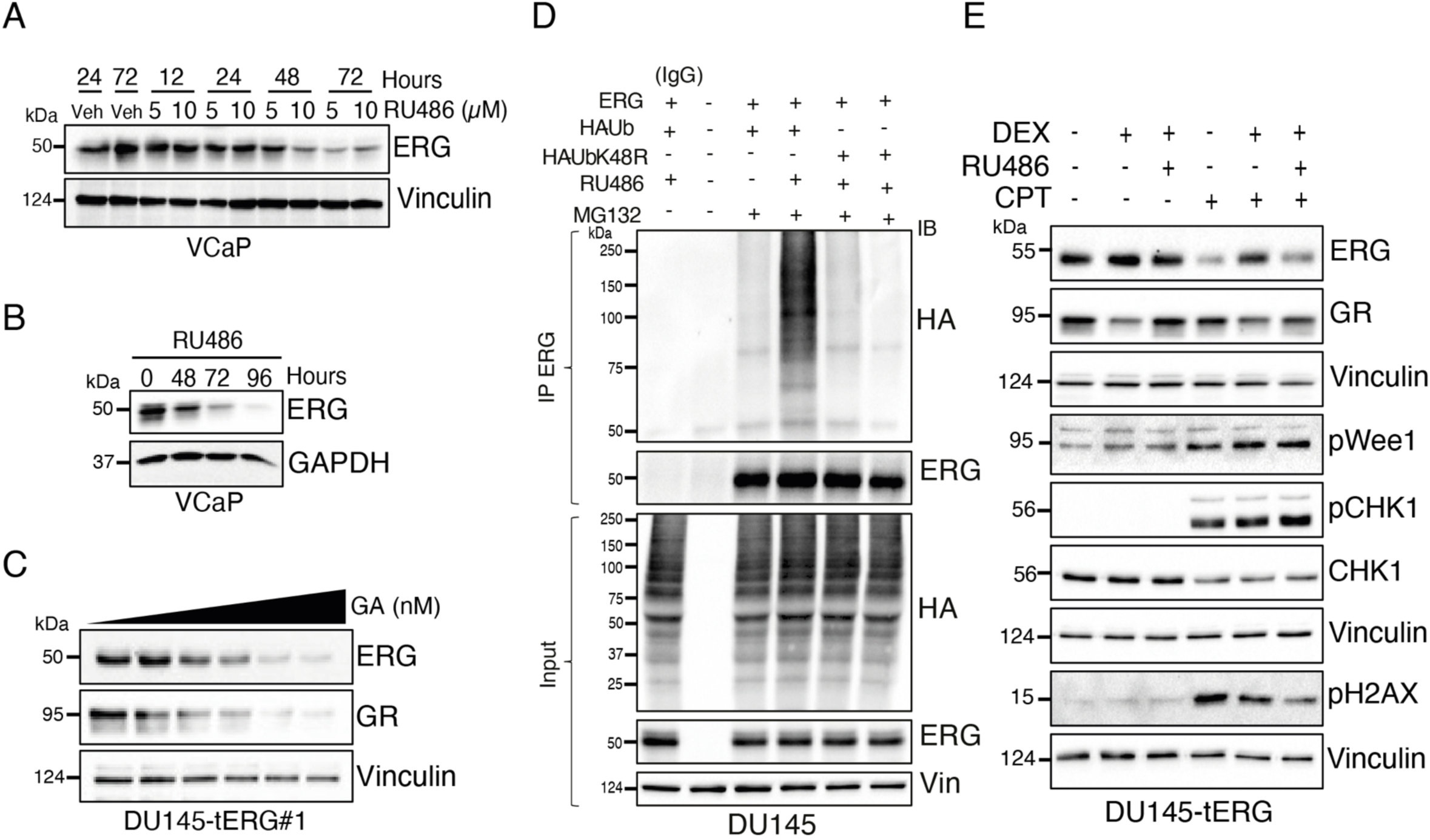
Inhibition of GR leads to ubiquitination and degradation of the ERG oncoprotein (see also Figure S3). (**A**) VCaP cells were treated for 12, 24, 48 and 72 hours with the indicated concentrations of RU486 (either 5 or 10 μM) and, thereafter, whole cell extracts were probed for the endogenous ERG or vinculin. Note that the control cells were treated with the vehicle (ethanol) for 24 or 72 hours. (**B**) VCaP cells were treated with RU486 (10 μM) for the indicated time intervals and ERG protein levels were assayed using immunoblotting. GAPDH served as a loading control. (**C**) DU145 cells stably overexpressing tERG were treated for 48 hours with increasing concentrations of GA. Post treatment, the cells were collected and their cleared extracts subjected to immunoblotting that used the indicated antibodies. (**D**) DU145 cells were transfected with HA-Ub and ERG encoding plasmids. Sixteen hours post transfection, the cells were treated for 48 hours with RU486. MG132 (25 μM) was added 8 hours prior to the end of the incubation. Cell extracts were subjected to immunoprecipitation using an anti-ERG antibody. Immunoblotting (IB) used an anti-HA antibody. A fraction (5%) of the total extract was used as the input control. (**E**) DU145-tERG cells were treated for 8 hours with camptothecin (CPT; 0.5 μM), in the absence or presence of DEX (1 μM), either alone or in combination with RU486 (10 μM). Cleared cell extracts were resolved using electrophoresis and western blotting that employed the indicated antibodies, including antibodies specific to the phosphorylated forms of CHK1, Wee1 and histone 2AX.

GR functionally depends on the HSP90 molecular chaperone ^45^. Because our results raised the possibility that GR can chaperone ERG, inhibiting the GR-HSP90 complex offers an alternative way to destabilize ERG using geldanamycin (GA), a specific HSP90 inhibitor. In agreement with this scenario, treatment of VCaP cells with GA decreased the levels of GR, and this was associated with decreased tERG levels (Fig. S3D). In addition, combining GA and RU486 further reduced tERG levels. According to an alternative interpretation, the previously reported ability of HSP90 to chaperone AR ^46^ may explain why GA could reduce the abundance of ERG. To examine this alternative model, we examined DU145-tERG cells, which express hardly detectable AR ^47^. Nevertheless, in these cells GA clearly induced degradation of both GR and ERG (Fig. 3C). In summary, inhibiting GR using either RU486 or GA decreased the abundance of tERG, in line with direct chaperoning of tERG by GR.

Our results predicted that blocking GR sorts the physically associated ERG proteins to intracellular degradation by means of a preparatory step involving conjugation of poly-ubiquitin chains that use lysine-48 of ubiquitin as a branching point. This model was tested in cells co-transfected with HA peptide-tagged ubiquitin constructs, either WT or the K48R mutant form of ubiquitin, along with a tERG plasmid. As shown in Figure 3D, immunoprecipitation detected a fraction of tERG that underwent poly-ubiquitination under basal conditions, and this relatively small fraction was strongly enhanced following treatment with RU486 and a 26S proteasome inhibitor (MG132). As expected, under the same conditions we detected no conjugation of the K48R ubiquitin mutant. In conclusion, HSP90-stabilized GR molecules physically bind with tERG and stabilize the bound oncoprotein. Consequently, inhibiting either GR, using RU486, or HSP90, using GA, shortens the half-life of ERG. In addition, we found that inhibiting GR instigates K48-branched chains of ubiquitin, which sort tERG molecules to degradation by the 26S proteasome.

It has been reported that irradiation or treatment with certain chemotherapeutic agents, such as camptothectin (CPT), enhances proteasomal degradation of both wild-type ERG and tERG in a mechanism that requires pre-phosphorylation of ERG at two sites: threonine-187 (by GSK3β) and tyrosine-190, by a cascade that includes ATR, CHK1 and WEE1 ^44^. We confirmed that CPT-induced DNA damage activated CHK1 and WEE1 in CPT-treated DU145-tERG cells and showed that this culminated in phosphorylation of histone H2AX (gamma-H2AX), a marker of double strand breaks (Fig. 3E). In line with the previous report, CPT enhanced ERG degradation. Still, treatment with DEX prevented stress-induced degradation of tERG, and as expected, RU486 nullified the effect of DEX in cells exposed to CPT. In summary, the GR-HSP90 complex not only physically binds with ERG but it also protects the complex from proteasomal degradation in both naive and chemotherapy-treated PCa cells. These observations might bear clinical significance for patients with tERG-positive patients, who receive a combination of chemotherapy and DEX.

### Tethering enables GR to augment the transcriptional activity of tERG

Upon hormonal stimulation, GR translocates to the nucleus to transcriptionally activate target genes containing the glucocorticoid response element (GRE). Alternatively, without binding directly to DNA, GR physically associates with and control tethered TFs, either coactivators or corepressors ^48^. To examine the prediction that GR enhances rather than represses the transcriptional program of ERG, we employed an ETS binding site (EBS)-luciferase reporter ^49^. As expected, when the reporter was expressed together with increasing amounts of an ERG plasmid, we observed gradually increasing luciferase signals (Fig. 4A). This signal was further increased when GR was co-expressed (Fig. 4B). To exclude the possibility that GR directly activates transcription from the EBS, the reporter was used in the absence of an ectopic ERG. As shown in Fig. S4A, GR alone could not enhance the luciferase signal in the absence of an exogenously expressed ERG, implying a piggyback (tethering) mode of GR action (see a scheme in Fig. 4B). Next, we confirmed that the GR-induced transcriptional activation of ERG could be modulated by GR-specific drugs: following co-transfection of an ERG plasmid and the EBS reporter, HEK293 cells were stimulated with DEX, RU486 or the combination. As anticipated, DEX robustly increased the EBS transcriptional signal, whereas the combination with RU486 abolished the effect of DEX (Fig 4C). In conclusion, the ability of ERG to up-regulate transcription from the EBS can be significantly enhanced by the agonist-activated form of GR.

**Figure 4:**
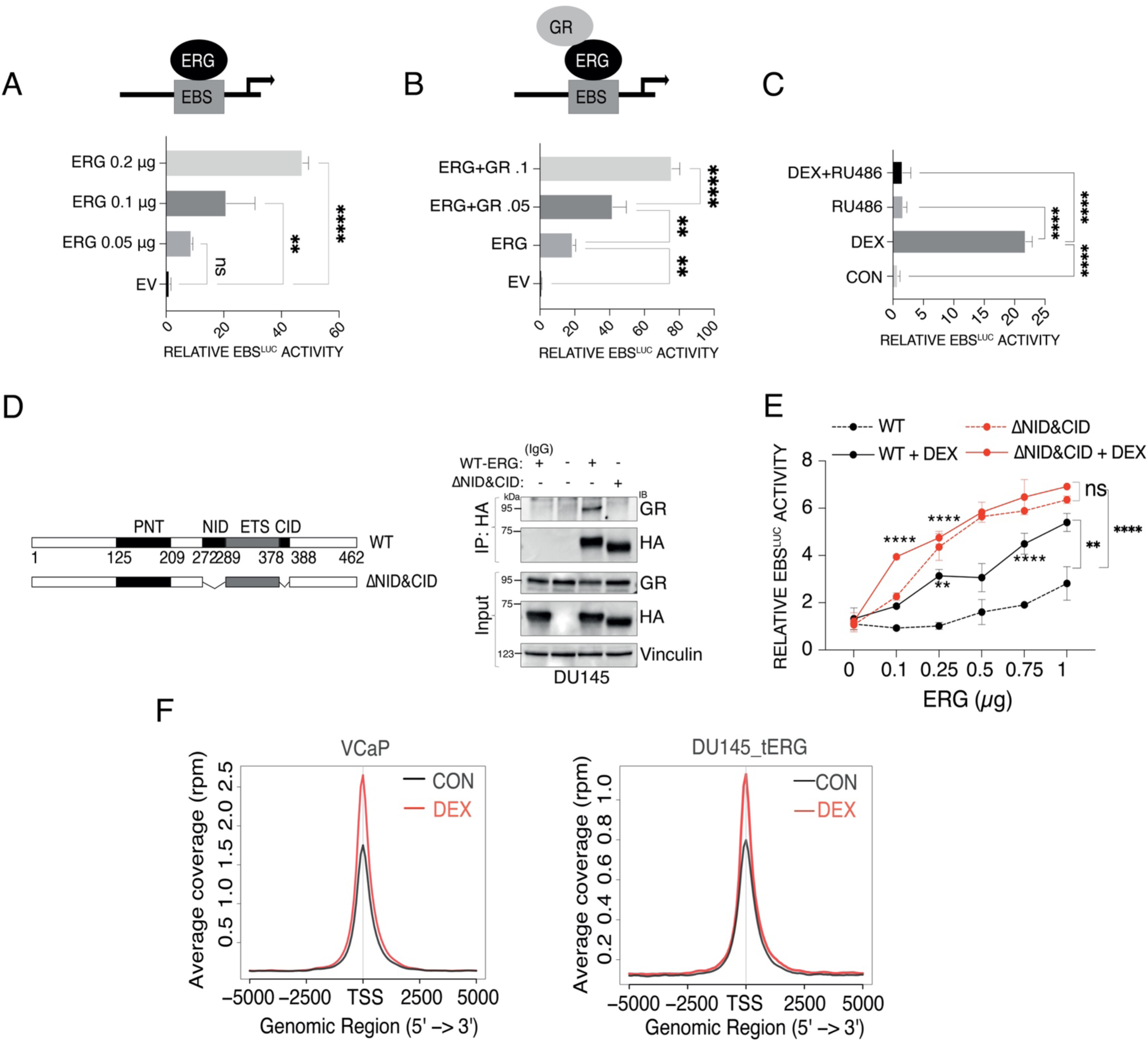
By relieving allosteric autoinhibition, GR enhances the transcriptional function of ERG (see also Figure S4). (**A**) HEK293T cells were co-transfected with an ETS binding site (EBS)-luciferase reporter plasmid and the indicated amounts of a plasmid encoding ERG. Luciferase activity was determined 24 hours later, using the Dual Luciferase Assay kit (from Promega). (**B**) HEK293T cells were co-transfected as in A with an EBS-luciferase reporter, an ERG plasmid and the indicated amounts of a plasmid encoding GR. Twenty-four hours later, we determined luciferase activity. (**C**) HEK293T cells growing in medium supplemented with charcoal-stripped serum (10%) were co-transfected with plasmids encoding EBS-luciferase and an ERG plasmid (or an empty vector). Sixteen hours post transfection, cells were stimulated for 60 minutes with DEX (1 μM), RU486 (1 μM) or the combination. Luciferase signals and p-values were determined. (**D**) (Left panel) Schematic representations of the domain structures of wildtype ERG, along with the corresponding double deletion mutant lacking both the N- and C- terminal autoinhibitory domains (ΔNID&CID). (Right panel) DU145 cells were transfected with plasmids encoding HA-tagged forms of ERG-WT and ERG-ΔNID&CID. Whole cell extracts were prepared twenty-four hours post transfection and processed for gel electrophoresis, or for a prior immunoprecipitation step that used an anti-HA antibody. Gel-resolved proteins were immunoblotted for GR, HA or vinculin (the loading control). IgG, control immunoglobulin. (**E**) HEK293T cells grown in medium supplemented with charcoal-stripped serum (10%) were co-transfected with an EBS-luciferase reporter, as well as with the indicated WT or mutant ERG plasmids at increasing doses. Sixteen hours post transfection, cells were stimulated with DEX (1 μM). Luciferase activity was determined using a kit. (**F**) CSS starved VCaP and DU145-tERG cells were treated for 1 hour, in duplicates, with either vehicle or DEX. Thereafter, cells were fixed and ChIP was performed using an antibody specific to ERG. The ChIP peak profile plots present genome-wide changes in ERG enrichment (average coverage) before (black) and after (red) stimulation with DEX. TSS, transcription start site.

To further support this scheme, we establish GR-knockout VCaP cells (Fig. S4B). Wild-type and KO cells were co-transfected with the EBS reporter, and luciferase signals were subsequently determined. In comparison to WT cells, GR-KO clones displayed relatively low EBS signals (Fig. S4C), but transfection of GR partly rescued the luciferase signal, in line with the proposed model. Since GR and ERG form a physical complex and each component of the complex binds with distinct DNA response elements, we next asked if ERG would trans-activate, or trans-repress, transcription from the GRE. To examine this, we transfected a GRE luciferase reporter, along with increasing amounts of an ERG plasmid. As expected, control experiments using an ectopically expressed GR showed enhanced transcription from the GRE (Fig. S4D). Interestingly, however, the ectopically expressed ERG inhibited, rather than enhanced basal expression from the GRE. This unexpected inhibitory effect was confirmed using another experiment that detected consistently reduced effects of GR on transcription from the GRE when ERG was overexpressed (Fig. S4E). Conceivably, opposite to the stimulatory effect of GR on transcription from the EBS, ERG reciprocally induces an inhibitory effect on transcription from the GRE.

### Transactivation by GR involves alleviation of an ERG allosteric inhibitory mechanism and enables GR to enhance transcription from the ERG’s DNA binding site

Previous analyses uncovered an intrinsic allosteric mechanism that autoinhibits ERG’s binding to DNA ^50^. Accordingly, two short segments of ERG, the N- and C-terminal inhibitory domains (NID and CID, respectively), each flanking the DNA binding region (ETS; see scheme in Fig. 4D), regulate intramolecular interactions. Our finding that GR engages the DNA binding domain of ERG, raised the possibility that GR can alleviate autoinhibition when in complex with ERG. To test this model, we utilized an HA-tagged double deletion mutant of ERG, ΔNID&CID, lacking both autoinhibitory domains. Control immunoprecipitation assays indicated that the double deletion mutant lost the ability to bind with GR (Fig. 4D). Next, increasing doses of plasmids encoding the mutant or the wild-type forms of ERG were expressed in hormone-starved cells, along with the EBS-luciferase reporter (Fig. 4E). As predicted, treatment of WT-ERG expressing cells with DEX was associated with increasing luciferase signals. In contrast, although the signals observed with the ΔNID&CID double mutant were higher, cell treatment with DEX did not further increase transcription. These observations are consistent with the autoinhibition model and they support the possibility that GR can physically or allosterically abolish ERG’s autoinhibition.

After demonstrating that GR can activate transcription from the EBS, we proceeded to use chromatin immunoprecipitation (ChIP) to investigate the genome-wide impact of GR on tERG’s chromatin-binding capacity. Before conducting the ChIP analysis, VCaP and DU145-tERG cells cultured in CSS (charcoal-stripped serum) were treated with DEX. In line with the promoter reporter assay findings, DEX stimulation led to approximately a 60% increase in the average ERG ChIP-seq signal in both VCaP and DU145-tERG cells (Fig. 4F). Collectively, the data in Figures 4 and S4 reveal mechanisms involving allosteric interactions and molecular tethering that likely enable GR to co-occupy tERG’s target promoters in patients expressing fused ETS genes.

### Inhibition of GR selectively decreases growth and survival of *TMPRSS2-ERG* positive cells

tERG-overexpressing PCa cells often rely on the fused gene for survival and proliferation ^51,52^. However, the relative contribution of the GR-ERG protein complex remains unknown. Hence, we firstly assayed ERG and GR levels in a panel of PCa cell lines (Fig. S5A), and then treated with RU486 a partly overlapping panel, which included RWPE, a normal prostate epithelial line, the tERG-expressing VCaP cells, PC3 cells, which express ETV4, and the tERG-low DU145 cells. The results showed that unlike the tERG-low/negative lines (DU145 and RWPE-1), the tERG-positive cells were sensitive to RU486 (Fig. 5A). Similarly, applying non-steroidal GR antagonists, instead of RU486, confirmed inhibition of the ETS-positive line, VCaP, but DU145 cells were not affected (Fig. 5B). In addition, we asked if inhibition of GR using RNA interference would similarly retard cell proliferation. GR was downregulated in VCaP, PC3, as well as in the tERG-low DU145 cells, using GR-specific siRNAs (Fig. S5B). This led to significant inhibition of the ability of VCaP and PC3 cells to incorporate a radioactive nucleoside into chromosomal DNA during mitosis (Fig. S5C). In contrast, we observed no significant inhibitory effect when DU145 cells (tERG-low) were treated with GR-specific siRNAs. Hence, we concluded that both pharmacological and genetic inhibition of GR can lead to reduced growth of ETS-positive PCa cells. Similar conclusions were reached on the basis of colony formation assays performed with VCaP, DU145 and PC3 cells (Figs. 5C and S5D). In line with these observations, we noted that treatment with RU486, or with non-steroidal GR antagonists, increased to variable extent both early and late apoptosis of ETS-positive cells (Figs. 5D and S5E). In similarity to the other assays, when tested on DU145 cells, RU486 induced no appreciable apoptosis. Next, we applied gene ablation and RNA interference as alternative strategies to address the function of the GR-ETS complex: in comparison to wildtype cells, GR-knockout VCaP cells displayed lower cell viability (Figs. S5F and S5G), DNA synthesis (Fig. S5H) and proliferation (Fig. S5I). Likewise, targeting ERG using specific siRNA nucleotides revealed that downregulation of *tERG* abolished sensitivity to RU486 (Figs. S5J and S5K). In conclusion, inhibition of either component of the GR-tERG complex can impair growth of several different PCa cells, implying that a substantial fraction of the mitogenic activity of tERG is attributable to the GR-tERG complex.

**Figure 5:**
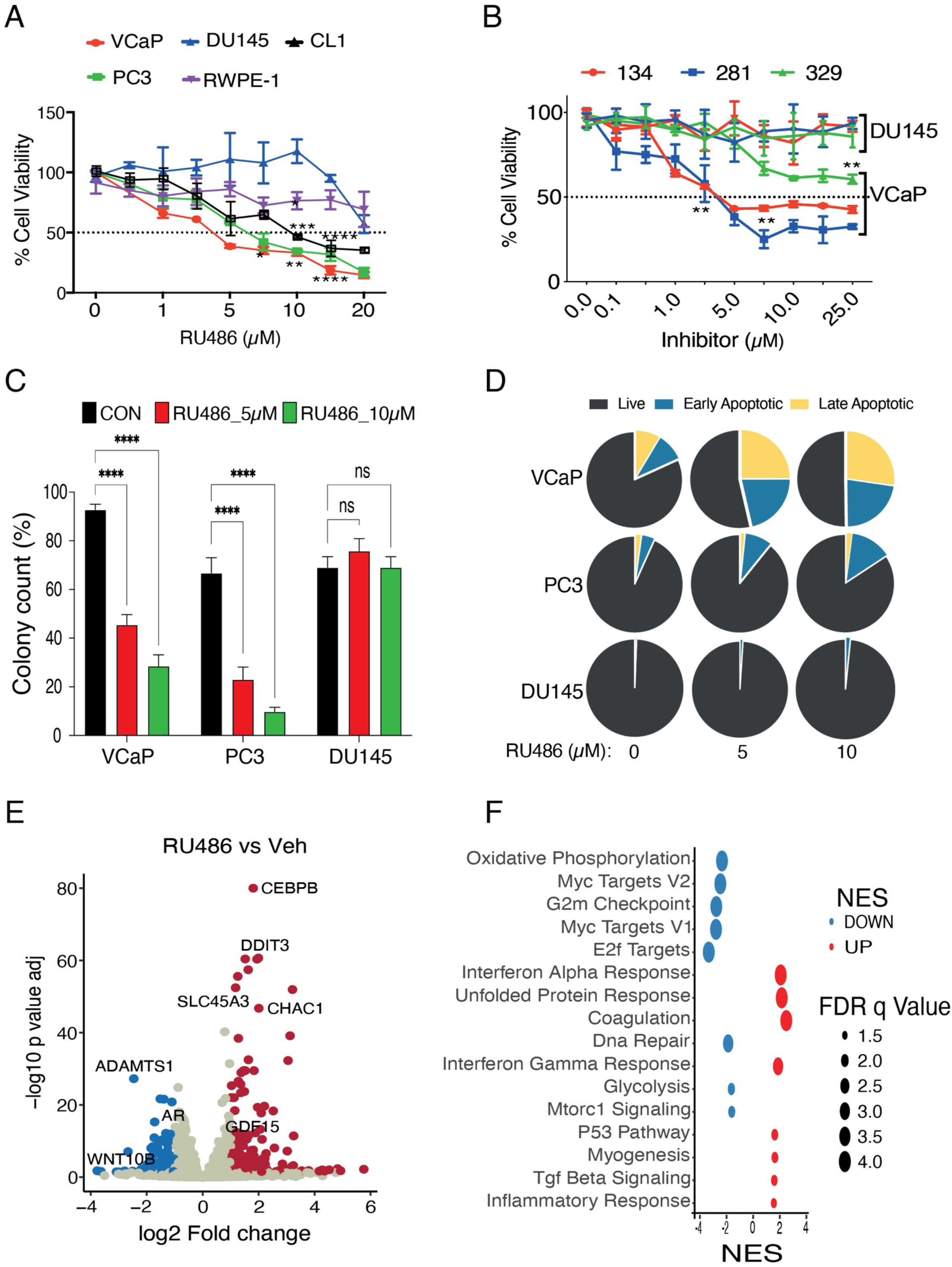
Inhibition of GR specifically decreases growth and survival of *tERG*-positive PCa cells (see also Figure S5). (**A**) The XTT cell viability assay was performed with the following prostate cell lines: VCaP (ERG+), PC3 (ETV4+), CL1 (ETV1+), DU145 (ERG-low) and RWPE-1 (normal prostate epithelial cells). All cell lines were treated in triplicates for 96 hours with increasing concentrations of RU486. (**B**) VCaP and DU145 cells were treated for 96 hours with increasing concentrations of the indicated non-steroidal inhibitors of GR, and later subjected to cell viability assays, as in A. (**C**) VCaP, PC3 and DU145 cells were sparsely seeded in 6-well plates and later untreated or treated on every other day with either vehicle or RU486 (5 or 10 µM). Ten days later, all cells were fixed and stained with crystal violet. The bar plots present quantification of colonies observed in 5 non-overlapping microscope fields. (**D**) The indicated cell lines were seeded in 100-mm dishes. Thereafter, they were treated for 72 hours with the vehicle (*CON*) or RU486 (1 μM). Shown are results of an apoptosis assay performed using an annexin V/7-AAD kit (from BioLegend). (**E**) VCaP cells were treated for 24 hours with RU486 (1 μM) or with the corresponding vehicle, and RNA was extracted for RNA-seq analysis. The volcano plot shows the differentially expressed genes. Significantly up-or down-regulated gens are colored in red and blue, respectively. The ERG target genes and the enriched apoptotic pathway genes are marked. (**F**) The presented pathway enrichment analysis of differentially expressed genes (DEGs) used EnrichR and selected genes with a fold change >1, or < -1, and an adjusted p-value smaller than 0.05. The most significantly enriched pathways from MSigDB Hallmark 2020 are depicted. FDR, false discovery rate.

To uncover transcriptional programs that may underlie the action of the GR-tERG complex, we treated VCaP cells with RU486, performed RNA-seq analysis (Fig 5E) and listed the pathways regulated by the differentially expressed genes (DEGs; Fig. 5F). This analysis selected the most significantly enriched pathways according to MSigDB Hallmark 2020 ^53^. As expected, this analysis identified previously reported ERG target genes, which were downregulated, along with apoptosis pathway genes that were up-regulated by RU486. For example, C/EBP-beta, which mediates expression of several interferon gamma regulated genes was upregulated in RU486-treated cells. Conversely, this drug inhibited expression of amphiregulin, a growth factor, and WNT10B, a canonical WNT ligand that controls malignant propensity. Likewise, we observed downregulation of genes regulating cell cycle progression (i.e., the E2F pathway), along with several *MYC* targets. The latter, as well as AR, FOXA1 and ERG, drive PCa cell proliferation ^54^. Taken together, these observations imply that GR cooperates with ERG by augmenting cell proliferation and overcoming apoptosis, while harnesseing the major TFs of PCa.

### Antagonizing GR, in similarity to lowering cortisol levels, selectively inhibits *TMPRSS2-ERG-*positive cell lines, as well as xenograft models

To examine the translational significance of the direct GR-to-tERG interactions, we treated three different PCa xenografts with either non-steroidal GR antagonists or with two clinically approved drugs: RU486 (mifepristone) and metyrapone (a steroidogenesis inhibitor that reduces cortisol levels ^55^). The following cells were implanted in immunocompromised mice: VCaP, PC3, and three derivatives of the tERG-low DU145 cells (parental, empty vector, EV, control cells, and tERG-overexpressing cells). Treatments making use of RU486, metyrapone and the non-steroidal compounds, especially CORT125329, clearly inhibited growth of the tERG-positive VCaP xenografts (Figs. 6A and S6A). In addition, PC3 cells that naturally overexpress another ETS protein, ETV4, were similarly inhibited by RU486 (Fig. S6B). In contrast, neither RU486 nor metyrapone inhibited growth of the parental DU145 cells (Fig. 6B). These observations led us to test in vitro the drug sensitivities of two clones of tERG-overexpressing DU145 cells. As expected, treatment with RU486 significantly reduced colony formation by the tERG-overexpressing clones, but growth of control cells pre-transfected with an empty vector (EV) was not inhibited by the drug (Figs. S6C and S6D). Consistent with the in vitro data, both the parental and the control DU145-EV cells displayed no response to RU486, but a tERG-overexpressing clone was significantly inhibited when tested in vivo (Figs. 6C and 6D). In summary, blocking GR signaling using two distinct strategies differentially inhibited growth of tERG-positive PCa cells, but the tERG-low DU145 cells displayed lack of responses.

**Figure 6:**
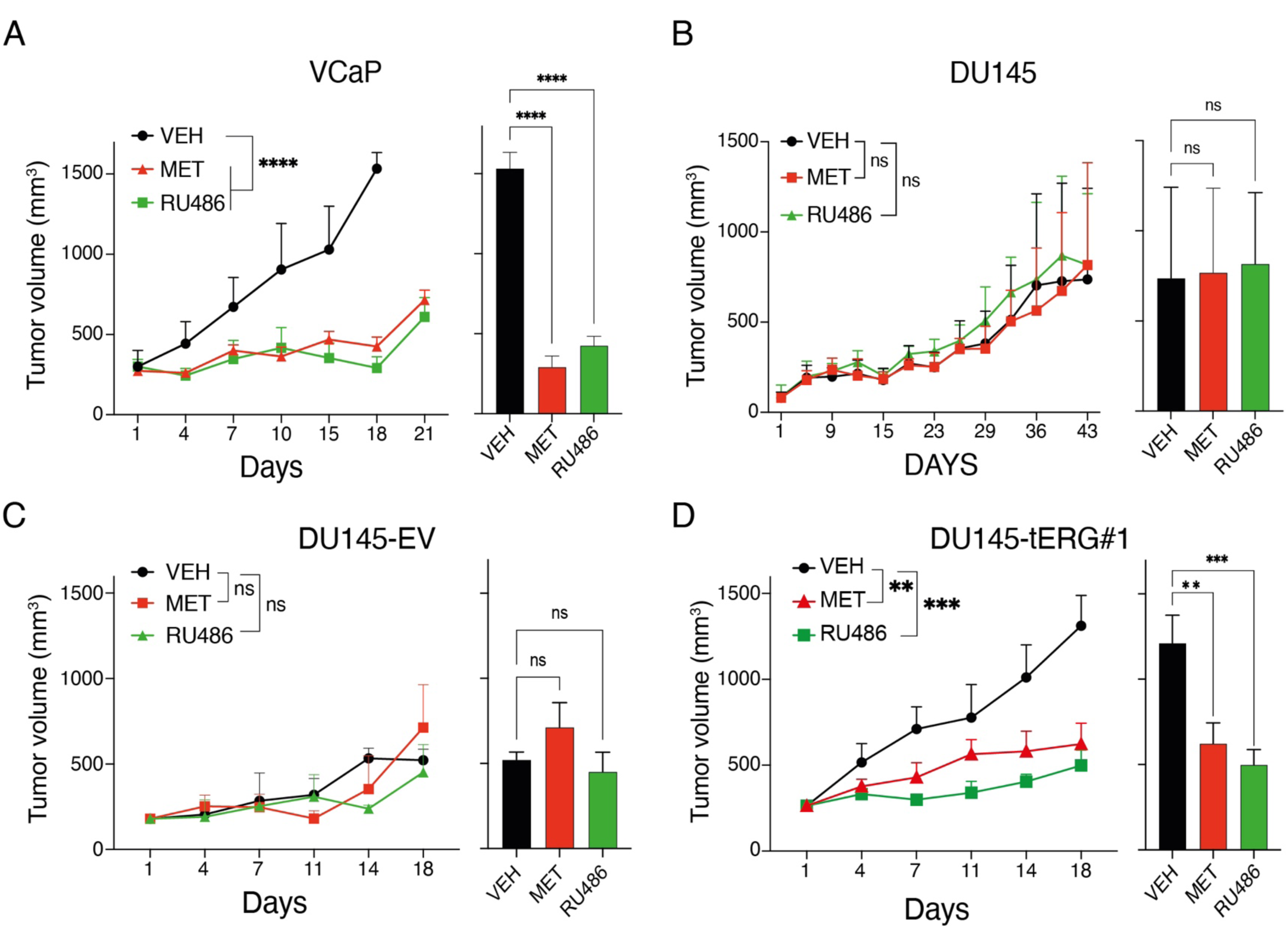
Both antagonizing GR and lowering cortisol inhibit *TMPRSS2-ERG^+^* xenograft models of PCa (see also Figure S6). (**A** and **B**) VCaP (2X10^6^) and DU145 cells (5X10^6^) were implanted subcutaneously in athymic mice. Once tumors became palpable, animals were randomized into three groups (5-6 animals per group), which were treated daily with vehicle, RU486 (1 mg/kg) or metyrapone (25 mg/kg). The rates of tumor growth are presented (left panels), along with the average tumor volumes on day 21 (VCaP) or day 43 (DU145), which are presented in the right panels. (**C** and **D**) DU145 cells stably expressing tERG (5X10^6^) and control DU145 cells (EV, 5X10^6^) were pre-established. Cells were implanted subcutaneously in nude mice and, once tumors became palpable, animals were randomized into three groups, which were treated daily with vehicle, RU486 (1 mg/kg), or with metyrapone (25 mg/kg). The bar graphs present tumor volumes corresponding to day 18. **, p < 0.01; ***, p < 0.001; ****, p<0.0001; ns, not significant.

### Combining GR and AR signaling blockers selectively inhibits tERG-positive xenografts derived from patients with PCa

In the next step, we examined in mice the potential benefit of combining GR inhibitors and enzalutamide. These experiments employed three patient-derived xenografts (PDX): two tERG-positive models (LuCaP 23.1 and LuCaP 35) and a tERG-negative PDX model (LuCaP 96) ^56^. Comparative analysis of the three PDX models confirmed very low ERG abundance in LuCaP 96 (Fig. S7A). Fragments of the LuCaP 23.1 model were implanted in NSG mice, which were later randomized into several groups that were treated with enzalutamide, RU486, or metyrapone. Daily treatments with enzalutamide, for 6 weeks, inhibited tumor growth, but a few weeks later all tumors adopted the rapid growth typical to the control untreated mice (Figs. 7A and S7B; see survival curves in Fig. 7B). Similarly, the RU486-treated group and the metyrapone-treated animals displayed delayed emergence of resistance. Importantly, however, combining enzalutamide and either GR-targeting drug resulted in monotonic, relatively slow tumor growth, which significantly prolonged animal survival. The other ERG-positive PDX model, LuCaP 35, displayed more aggressive growth and only weak response to enzalutamide monotherapy (Figs. 7C and S7C). Still, combining enzalutamide and either RU486 or metyrapone partly but significantly inhibited tumor growth, as well as prolonged animal survival (Figs. 7C and 7D). Interestingly, the efficacies achieved by each pair of drugs were comparable and significantly better than the effect of enzalutamide monotherapy. In conclusion, both ERG-positive PDX models we tested clearly displayed sensitivity to the doublet combinations of enzalutamide and an anti-GR drug.

**Figure 7:**
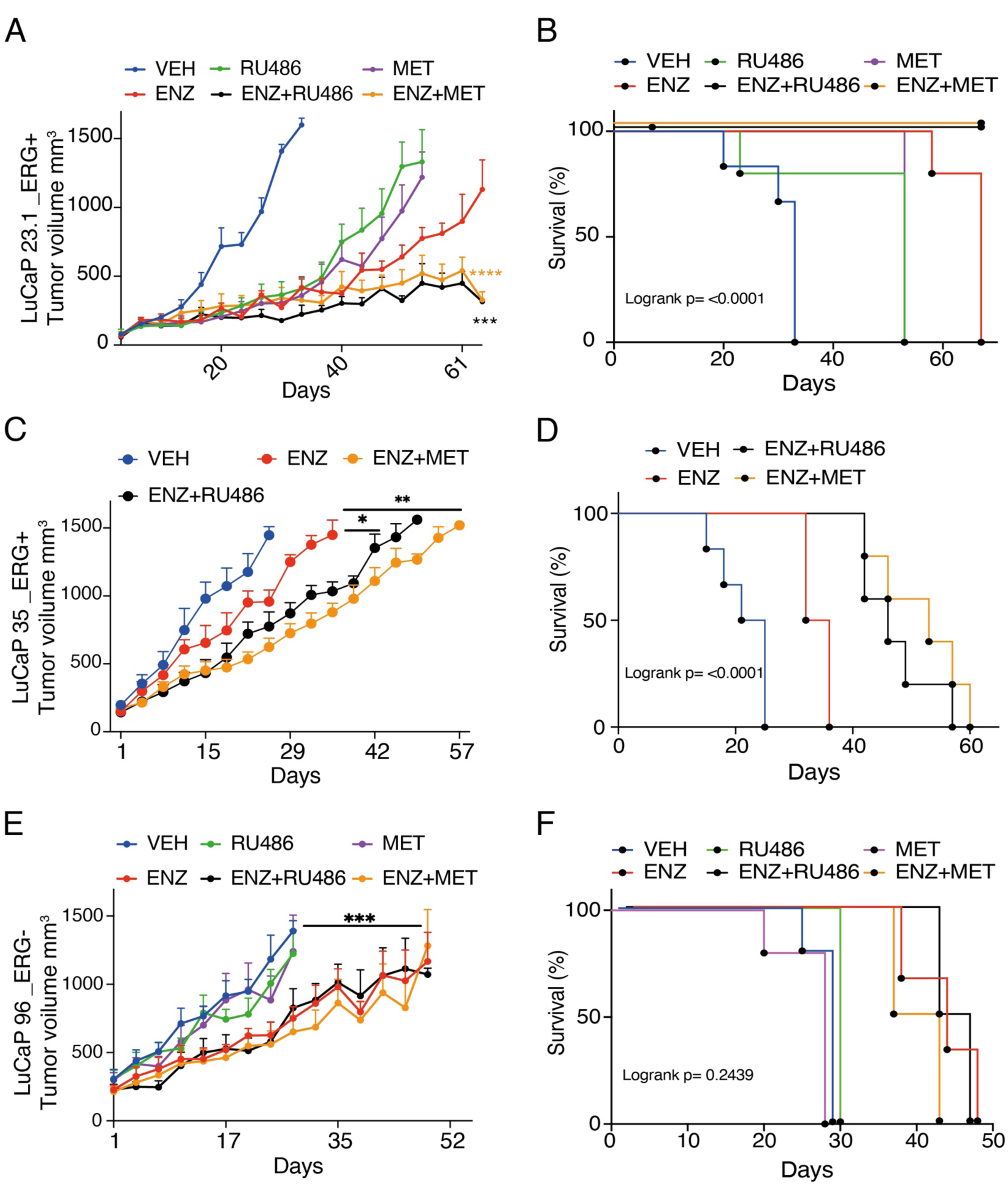
Combinations of GR and AR signaling inhibitors selectively retard tERG-positive xenografts derived from patients with PCa (see also Figure S7). (**A** and **B**) Fragments of the tERG-positive LuCaP 23.1 PDX model were subcutaneously implanted in the flanks of 4-6 NSG mice. Once tumors reached 150 mm^3^, animals were randomized into the indicated groups, which were daily treated with enzalutamide (ENZ; 10 mg/kg, oral gavage), RU486 (1 mg/kg, intraperitoneal injection), metyrapone (MET; 25 mg/kg, oral gavage), or with the indicated drug combinations. Tumor volumes were monitored twice per week and body weights were recorded once per week. Shown are average tumor growth rates of mice treated with single drugs or with the indicated dual drug combinations. Also shown are Kaplan-Meier plots corresponding to survival analyses. Note that mice were euthanized when tumor size reached 1,500 mm^3^. (**C** and **D**) Fragments of the tERG-positive LuCaP 35 PDX model were subcutaneously implanted in the flanks of NSG mice and treated as in A. The Kaplan-Meier plots present the respective animal survival data (D). (**E** and **F**) Fragments of the tERG-negative LuCaP 96 PDX model were subcutaneously implanted in the flanks of NSG mice and treated as in A. The Kaplan-Meier plots present the respective animal survival data (F). *, p < 0.05; **, p < 0.01; ***, p < 0.001; ****, p<0.0001; ns, not significant.

As expected from the in vitro data and from the results obtained with xenografts of the ERG-low DU145 cell line, the third PDX model, LuCaP 96, which is tERG-negative and expresses only low ERG levels, displayed resistance to anti-GR drugs (Figs. 7E-F and S7D). Although this model partly responded to enzalutamide, neither anti-GR drug could enhance enzalutamide’s effect in terms of tumor growth and animal survival. Hence, in view of the herein reported physical interaction between ERG and GR, the observations made in tumor-bearing animals indicate that simultaneously targeting both component of the GR-tERG complex has an added pharmacological value that is limited to fusion-positive tumors.

In conclusion, analysis of patient data from two clinical trials suggested that the presence of *TMPRSS2-ERG* (tERG) favors resistance to ARSIs and initiates drug-induced up-regulation of GR. To resolve the underlying mechanism, we performed protein complementation assays and other tests. These studies revealed that the glucocorticoid receptor physically interacts with tERG, relieving its intrinsic auto-inhibition and protecting it from both basal and chemotherapy-induced degradation. As a result, GR boosts activation of tERG target genes, including the MYC proto-oncogene. Critically, this mechanism is absent in tERG-negative tumors, which are more responsive to ARSIs. Supporting the notion that GR-*tERG* complexes create specific vulnerabilities, we observed that inhibiting GR or reducing cortisol synthesis promoted apoptosis and suppressed growth in *tERG*-positive prostate cancer cells but had lesser effects on *tERG*-negative cells. In animal models with *tERG*-positive tumors, combining antiandrogens with GR signaling inhibitors demonstrated cooperative anti-tumor effects. Thus, by identifying GR as a critical oncogenic partner of tERG, this study uncovers promising therapeutic strategies to overcome drug resistance and improve outcomes for a significant subset of patients with PCa.

## Discussion

Two decades ago, it was discovered that approximately every other patient with PCa expresses fusions between *TMPRSS2* and an *ETS* family gene ^2^. However, unlike other fusion genes and activated mutant oncogenes like BCR-ABL and B-RAF, which are targetable by several anti-cancer drugs, no clinically approved drug targets TMPRSS2-ETS proteins, and the presence of the rearranged genes has not been translated to predictive biomarkers. The clinical data analyses we present confirmed bimodal distribution of untreated PCa tumors: those harboring *TMPRSS2-ERG* displayed much higher expression of ERG. Importantly, the respective group of patients exhibited statistically significant resistance to treatments that made use of antiandrogens. As far as we are aware, this is the first demonstration of the ability of *TMPRSS2-ERG* to confer resistance to antiandrogens. Because the resistant patients displayed relatively high GR levels and previous studies established critical roles for GR in resistance to antiandrogens ^20,57^, our search for the underlying mechanism focused on GR and its reported ability to form complexes with several ETS family members ^58^. Protein complementation and additional assays revealed that GR directly interacts with ERG, thereby it alleviates ERG’s allosteric autoinhibition, as well as stabilizes ERG and boosts transactivation of the respective target genes. Thus, although *TMPRSS2-ERG* gene fusions emerge at relatively early stages of PCa development ^6^ and they are associated with aggressive phenotypes, our results portray the respective tumors as vulnerable cancers due to acquired dependence on several onco-proteins: AR and GR in the first place, but also HSP90, which controls both receptors, and tERG, which is inducible by androgens. The dependence on GR becomes especially critical when the AR pathway is blocked: normally AR suppresses GR ^23^, but following treatment with antiandrogens GR undergoes upregulation and provides an escape route that engages the majority of the AR target genes. The inferred mechanism explains why our tERG-positive xenografts displayed sensitivity to combinations of GR and AR blockers, but the tERG-negative models we utilized were significantly less responsive.

Beyond the implications for resistance to antiandrogens and the therapeutic promise offered by combining GR and AR inhibitors, specifically when treating tERG-positive patients, our study is relevant to the application of corticosteroids in PCa treatment. Although most castration-resistant prostate cancer patients benefit from abiraterone plus prednisone, resistance eventually emerges. However, this may be reversed by switching to the longer lived corticosteroid, dexamethasone ^59–61^. Likewise, daily oral corticosteroids are routinely combined with abiraterone to mitigate symptoms of mineralocorticoids ^62^ and, similarly, prednisone augments the efficacy of docetaxel in patients with PCa ^63^. However, according to our observations, corticosteroids might accelerate disease progression, but this appears to be limited to tumors expressing a fused form of an ETS family member. In line with this scheme, high GR expression is associated with an increased risk of disease progression in several carcinomas ^64^ and treatment of castration resistant PCa cells with AR inhibitors frequently involves up-regulation of GR, which stimulates a subset of AR-regulated genes ^20,21^. In addition, the uncovered ability of hormone-activated GR molecules to bind with and enhance the oncogenic activity of tERG might explain the recently reported post hoc analysis of the AFFIRM phase III clinical trial (NCT00974311). This study randomized 1,199 patients to enzalutamide or placebo and allowed corticosteroids at study entry and during the trial. Along with confirming the clinical efficacy of enzalutamide, the new study addressed the clinical impact of concurrent corticosteroid use (CCU) on enzalutamide-treated patients with metastatic disease ^29^. Unexpectedly, a multivariate analysis found that baseline CCU was independently associated with decreased overall patient survival. Interestingly, an earlier post hoc analysis of another clinical trial, COU-AA-301, which tested abiraterone plus prednisone versus prednisone in metastatic PCa patients made a similar observation but reached a different conclusion ^28^: Although in similarity to the AFFIRM data, COU-AA-301’s data found that baseline corticosteroids were associated with shorter overall patient survival, the authors concluded that this inferiority was due to an association between the use of baseline corticosteroid and patients having worse disease characteristics. In aggregate, the reviewed lines of evidence raise the possibility that corticosteroid therapy may be contraindicated in PCa patients with tumors expressing an ETS fusion gene.

In conclusion, our in vivo and other observations identify a protein complex comprising GR and tERG as a driver of resistance to antiandrogens. In addition, our results shed light on the underlying molecular mechanism: blocking AR elevates GR, which directly boosts the oncogenic attributes of tERG. Thus, GR agonists like DEX and prednisone might accelerate progression of prostate tumors expressing tERG. From the medical point of view these molecular mechanisms may translate to a new predictive biomarker and a recommendation to limit the use of CCU to patients with tERG-negative PCa. Thus, the reported findings might motivate development of new anti-GR drugs, such as small molecules sharing the ability to intercalate into the newly identified GR-ERG cleft. Alternatively, if validated by additional studies, our observations might justify the development of next-generation GR antagonists, such as the non-steroidal CORT agents we tested herein, AR-GR dual antagonists and steroidal agents like ORIC-101 ^65^.

## Acknowledgments

We thank all members of our laboratories for kind help and insightful comments. We also thank Drs. Swati Srivastava for establishing PCA, Haim Werner (Tel Aviv University) for PC3 cells and Marie-Helene David (INSERM) for the EBS reporter. We are grateful to Drs. Johann de Bono and Charles Sawyers for helpful feedback and comments. YY is the incumbent of the Harold and Zelda Goldenberg Professorial Chair in Molecular Cell Biology. Our studies were supported by the Israel Science Foundation, European Research Council (ERC), the Israel Cancer Research Fund (ICRF) and the Dr. Miriam and Sheldon G. Adelson Medical Research Foundation. This research was supported in part by the Intramural Research Program of the National Institutes of Health, NCI. We thank Corcept for providing CORT125134, CORT125281 and CORT125329.

## Author contributions

Conceptualization, A.S. and Y.Y.; Methodology, M.Z., R.B-G.; Formal Analysis, D.R.G., Y.V., E.W., E.D., L.R., J.B.R., R.T.L.; Investigation, A.S, R.C., A.V., S.G., A.K., S.K., B.R.S., N.B.N., F.K., N.G., T.A.; Resources, Y.P., H.H. (H.Huang), W.M.W., E.C., H.H. (H. Hunt), A.E.G., A.G.S.; Writing – Original Draft, A.S. and Y.Y.; Writing – Review and Editing, all authors.; Supervision, A.A.S, R.O., A.A., W.L.D., A.G.S., S.L., E.R., I.A., Y.Y.; Project Administration, A.S. and Y.Y.; Funding acquisition, E.R., Y.Y.

## Inclusion and Diversity

We support inclusive, diverse, and equitable conduct of research.

## Materials and methods

### Clinical samples and patient data analysis

The following clinical samples and datasets were analyzed in this study: (i) Patient samples and patient data from clinical trial NCT02430480 ^26,27^ were available to us. This clinical study recruited 37 men with PCa who received ADT and enzalutamide for 6 months. Biopsy samples were used to derive immunohistochemical data, whole-exome and RNA-sequencing information. We performed additional IHC staining of baseline and post-treatment biopsies (see Figure S1). (ii) Similarly, we reanalyzed clinical data from NCT01576172 ^30^. All patients of trial underwent metastatic site biopsy and assigned to either abiraterone plus prednisone, or to a combination of a PARP-1 inhibitor, veliparib and abiraterone plus prednisone. We only analyzed patients who were untreated with veliparib (58 patients). (iii) A dataset colllected by the West Coast Dream Team collaborative group, who performed whole genome and RNA sequencing of tumor biopsies from patients with metastatic castration-resistant prostate cancer^33^.

### Cell lines and reagents

PC3 cells were received from Prof. Haim Werner (Tel Aviv University). Other cells were from the American Type Tissue Culture Collection (ATCC). Human embryonic kidney cells, HEK293T, were cultured in Dulbecco’s modified Eagle (DME) medium supplemented with fetal bovine serum (FBS; 10%). DU145 and PC3 cells were grown in Roswell Park Memorial Institute (RPMI) medium supplemented with 10% FBS. VCaP cells were grown in RPMI supplemented with 10% FBS, glutamine and sodium pyruvate. siRNA transfections used ON-Target SMART oligonucleotides from Dharmacon.

### Generation of stable cell derivatives

To knockout GR in VCaP cells, we used the CRISPR–Cas9 system and created a double-stranded break next to the Protospacer Adjacent Motif (PAM). The target site was selected using the ENSEMBL database. The selected targets (21 bp) included the PAM sequences in exon 5. For the establishment of DU145-tERG overexpressing cells, lentiviral particles were produced in HEK293T cells by co-transfecting lentiviral expression vectors containing the coding region of ERG (pLX304-ERG), together with second-generation viral packaging plasmids (VSVG; Addgene #14888) and psPAX2 (Addgene #12260). Twenty-four hours after transduction, the virus-containing medium was replaced with selection media containing blasticidin.

### Protein complementation assay (PCA)

The PCA method, which employs two inactive fragments of luciferase, was used essentially as previously described ^36^. Prior to the assay, HEK293T cells were transfected in 48-well tissue culture plates with the Gluc1 and Gluc2 plasmids (25 ng, each), using the JetPEI reagent. Twenty-four hours later, the cells were extracted in luciferase lysis buffer (25 mM Tris, pH 8.5, 150 mM NaBr, 5 mM EDTA, 0.1% NP40, 5% glycerol, 65 μM sodium oxalate, 0.5 mM reduced glutathione and 0.5 mM oxidized glutathione). Native coelenterazine (Nanolight) was diluted in luciferase assay buffer (25 mM Tris,pH 7.75, 1 mM EDTA, 0.5 mM reduced glutathione, 0.5 mM oxidized glutathione and 75 mM urea) to a final concentration of 20 μM. Luminescence signals were determined using a Veritas microplate luminometer (Turner BioSystems).

### Cell lysis, immunoblotting and co-immunoprecipitation assays

Post treatment, cells were washed twice with ice cold saline. Cell lysates were collected in a mild lysis buffer (50 mM HEPES, pH 7.5, 10% glycerol, 150 mM NaCl, 1% Triton X-100, 1 mM EDTA, 1 mM EGTA, 10 mM NaF and 30 mM β-glycerol phosphate). Proteins were immunoprecipitated from cell lysates using beads conjugated to an antibody. Following 2 hours of incubation at 4°C, complexes were washed three times and bound proteins were eluted in concentrated gel sample buffer. Eluates were subjected to electrophoresis and immunoblotting.

For immunoblotting, cleared cell lysates were resolved using electrophoresis, which was followed by electrophoretic transfer to a nitrocellulose membrane. Membranes were blocked in TBS-T (tris-buffered saline containing Tween-20) containing 1% low-fat milk, blotted overnight with a primary antibody, washed three times in TBS-T, incubated for 30 minutes with a secondary antibody linked to horseradish peroxidase, and washed once again with TBS-T. Immunoreactive bands were detected using the ECL reagent (Biorad, USA).

### Cell growth assays

For XTT assays, cells (2X10^4^) were seeded in a 96-well plate, in triplicates, and treated with drugs for the indicated time intervals. Subsequently, XTT was added and following 3 hours at 37°C we determined optical density at 490 and 640 nm. For the thymidine incorporation assay, cells were plated onto 24-well plates at a density of 5X10^4^ cells/well, followed by the indicated treatments. Sixteen hours later, serum-containing media were replaced with fresh serum-free medium containing ^3^[H]-thymidine (1 μCi). After 48 hours, the reaction was terminated by the addition of ice-cold trichloroacetic acid (5%; TCA). Five minutes later, the cells were solubilized in 1N NaOH (for 10 minutes) followed by 1N HCl. Samples were collected into scintillation vials containing scintillation fluid. Radioactivity was determined in a scintillation counter. The results shown are representative of experiments performed in quadruplicates.

### Colony formation and apoptosis assays

Cells (150-300) were seeded in 6-well plates. Ten days later, cells were fixed in paraformaldehyde (4%) and stained with crystal violet. Cells were then photographed and analyzed using ImageJ. Apoptosis assays were performed using the FITC Annexin V Apoptosis Detection Kit with 7-AAD (from BioLegend) and analyzed using flow cytometry. The assays were performed on a BD FACSAria Fusion instrument controlled by BD FACS Diva software v8.0.1 (BD Biosciences). Further analysis was performed using the FlowJo software v10.2 (Tree Star).

### RNA isolation and real-time PCR analysis

Total RNA was extracted using the PerfectPure RNA Cultured Cell Kit (5-prime, Hamburg) according to the manufacturer’s instructions. RNA quantity and quality were determined using the NanoDrop ND-1000 spectrophotometer (Thermo Fischer Scientific, Waltham, MA). Complementary DNA was synthesized using the High-Capacity Reverse Transcription kit (Applied Biosystems, Life Technologies, Carlsbad, CA, USA). Real-time qPCR analysis was performed with SYBR Green (Applied Biosystems) and specific primers on the StepOne Plus Real-Time PCR system (Applied Biosystems). qPCR signals (cT) were normalized to beta2-microglobulin (B2M).

### Luciferase reporter assays

Cells were co-transfected with a luciferase plasmid containing the consensus glucocorticoid response element (GRE) or the ERG’s DNA binding site (EBS). Additionally, the pGL3-Control vector encoding Renilla luciferase (Promega, Madison, WI) was transfected as a control for transfection efficiency. Luciferase activity was determined using the Dual-Luciferase reporter assay system, according to the manufacturer’s instructions (Promega). Firefly luciferase luminescence values were normalized to Renilla luminescence and quantified relative to control.

### Immunofluorescence analyses

Cells were washed with saline and treated with an acidic buffer (100 mM glycine-HCl, pH 3.0) for 3 minutes. Later, cells were washed and fixed for 15 minutes using 4% PFA. Next, cells were washed and permeabilized for 10 minutes with 0.2% Triton X-100 in saline, and then subjected to blocking using 3% albumin. Thereafter, the cells were incubated overnight with primary antibodies specific to ERG or GR. After incubation, cells were washed and incubated with a secondary antibody (from Thermo Fisher Scientific) and DAPI, followed by washing and mounting on slides for imaging.

### Ubiquitination assays

Cells were co-transfected with HA-tagged ERG and Flag-tagged wildtype and K48R ubiquitin plasmids. Post-transfection, cells were treated for 48 hours with RU486. MG132 was added 8 hours prior to the end of the incubation. Ubiquitinated ERG from cell lysates was immunoprecipitated using an anti-HA antibody. Immunoblotting used an anti-FLAG antibody. A fraction (5%) of the total extract was used as the input control.

### Subcellular fractionation

Cell pellets were lysed in 0.1 ml cytoplasmic lysis buffer (10 mM HEPES pH 7.9, 10 mM KCl, 0.1 mM EGTA, 0.1 mM EDTA, 1 mM DTT and 0.5% NP-40). The cytoplasmic fraction was collected using centrifugation (600 g for 5 minutes). Nuclei were washed and resuspended in 50 µl nuclear lysis buffer (20 mM HEPES pH 7.9, 0.4 M NaCl, 1 mM EGTA, 1 mM EDTA and 1mM DTT) using repeated freezing and thawing. Supernatants containing the nuclear fraction were collected by centrifugation at 12,000 rpm for 20 minutes.

### Nucleotide sequencing of RNA

RNA was isolated using Dynabeads mRNA Direct Kit (Thermo FisherScientific). NGS libraries were prepared using a modified version of Transeq. In brief, RNA was barcoded and reverse-transcribed using poly-T primers, followed by the addition of an exonuclease to remove excess of the PCR primers. The single-stranded cDNA was converted to a double-stranded DNA. The template DNA was then removed using DNase, and the generated RNA was fragmented and ligated to barcoded Illumina adapters. Reverse transcription of the ligation product was performed using primers specific to the Illumina’s adapters, and libraries of the resulting cDNA were generated and enriched by performing 12–15 PCR cycles. RNA-seq libraries (pooled at equimolar concentrations) were sequenced on an Illumina NextSeq 500 at a median sequencing depth of ∼10 million reads per sample. Sequences were mapped to the human genome (hg38), demultiplexed and filtered.

### Analysis of RNA sequencing data

Single-end 100 bp reads were sequenced on Illumina NovaSeq. We obtained ∼12 million reads per sample. Poly-A/T stretches and Illumina adapters were trimmed from the reads using cutadapt and reads shorter than 30 bp were discarded. Reads were mapped to the human reference genome GRCh38_p13 using STAR ^66^. Reads with the same UMI were removed using the PICARD MarkDuplicate tool. Gene expression levels were quantified using htseq-count. Differentially expressed genes were identified using DESeq2 ^67^ with the betaPrior, cooksCutoff and independentFiltering parameters set to False. Raw P values were adjusted for multiple testing using the procedure of Benjamini and Hochberg ^68^. Sneakmake was used to run the pipeline.

### Pathway analysis

The tool EnrichR ^69^ was used to perform pathway enrichment analysis. Selected genes were used for overrepresentation analysis; in the gene expression analysis, genes with a fold change of at least 1, or -1, and an adjusted p-value smaller than 0.05 were analyzed.

### Chromatin immunoprecipitation (ChIP)

Cell fixation, chromatin immunoprecipitation and nucleotide sequencing were performed as described ^70^. ChIP assays were carried out using approximately 10 million VCaP cells per sample. Cells were crosslinked for 10 minutes in 1% formaldehyde at 37°C. This reaction was subsequently quenched in 2.5 M glycine for 5 minutes. Chromatin from formaldehyde-fixed cells was fragmented to a size range of 200–700 bases using a sonicator. Solubilized chromatin was immunoprecipitated overnight at 4°C with GR-or ERG-specific antibodies. Antibody-chromatin complexes were pulled-down using protein G-Dynabeads (Life Technologies), washed, and then eluted. After crosslinking reversal, RNase A and proteinase K treatment, the immunoprecipitated DNA was extracted using AMP Pure beads (Beckman Coulter). During ChIP-seq library preparation, DNA isolated from ChIP experiments, as well as input control DNA, was end repaired, A-tailed, ligated to barcoded Illumina adaptors, PCR-amplified, and pooled for sequencing.

### Prostate cancer animal models

All animal experiments were approved by the Weizmann Institute’s Animal Care and Use Committee and the Institute’s Review Board (IRB). Male CB17/*SCID* mice (5-6 weeks old) were subcutaneously implanted in the right dorsal flank with 2.5 million VCaP, PC3 or DU145 cells suspended in saline (0.1 ml). Three prostate cancer PDX models, LuCaP 23.1 (ERG positive), LuCaP 35 (ERG positive) and LuCaP 96 (ERG negative), were previously established^71^. These models were expanded in NSG mice. Following euthanasia, tumors were removed from donor mice and cut into small fragments. A small pouch was made in the lower back of 5-6 weeks old mice and one tumor fragment was later inserted into the pouch. Mice were labelled with RF identification chips (from Trovan, Melton, UK). Tumor volume (*V*/mm^3^) was estimated using vernier caliper measurements of the longest axis, *α*/mm, and the perpendicular axis, *b*/mm. Tumor volume was calculated in accordance with the equation *V* = (4*π*/3) x (*α*/2)^2^ x (*b*/2). When the volume of xenografts reached approximately 150 mm^3^, mice were randomized into groups and treatments were initiated. Animals were intraperitoneally treated once per day with RU486 (1 mg/kg; in 0.1% Tween 80) or with Corcept compounds (50 mg/kg; in 0.1% Tween 80, 0.5% hydroxypropyl methylcellulose, HPMC). Alternatively, mice were orally treated with enzalutamide, (10 mg/kg; 0.1% Tween 80, 0.5% HPMC) or metyrapone (25 mg/kg, in water). Animals were euthanized when tumors reached 1500 mm^3^.

### Immunohistochemical analyses

Formalin-fixed, paraffin embedded blocks of matched untreated biopsy tissue or posttreatment whole-mount prostatectomies were sectioned at 5µm thickness onto SuperFrost Plus charged glass slides. Slides were stained with antibodies against ERG (clone EPR3864) and GR (clone D6H2L) and analyzed as previously described ^26^. Briefly, after deparaffinization and rehydration, antigen retrieval was performed in a pressure cooker at 110°C for 15 minutes in Diva Decloaker buffer (Biocare), and stained for ERG (1:200 dilution) or GR (1:400 dilution) for 1 h using a Biocare IP FLX autostainer. Secondary detection was performed with Mach 4 polymer/probe reagents, developed with DAB, and counterstained with hematoxylin. After mounting, slides were scanned using an AxioScan.Z1 slide scanner. Fully quantitative enumeration of tumor cells expressing ERG and GR was performed using Definiens XD 64 software with 12 staining subsets per slide and segmentation level 9. A percent positive weighted histology index (between 0-1) was used to report expression of ERG and GR nuclear intensities. Semi-quantitative H-scoring (between 0-300) was used to report the nuclear intensity of ERG and GR in prostatectomy tissues. All tissue analyses were performed by a genitourinary pathologist.

### Statistical analyses

All data were analyzed using the Prism Graphpad software and R. Statistical analyses were performed using t-test and one-or two-way ANOVA with Tukey’s or Dunnett’s tests (*, p≤0.05; **, p≤0.01; ***, p≤0.001; ****, p≤0.0001).

## References

1. Cancer Genome Atlas Research, N. The Molecular Taxonomy of Primary Prostate Cancer. Cell 163, 1011–25 (2015).

2. Tomlins, S.A. et al. Recurrent fusion of TMPRSS2 and ETS transcription factor genes in prostate cancer. Science 310, 644–8 (2005).

3. Grasso, C.S. et al. The mutational landscape of lethal castration-resistant prostate cancer. Nature 487, 239–43 (2012).

4. Robinson, D. et al. Integrative clinical genomics of advanced prostate cancer. Cell 161, 1215–1228 (2015).

5. Paulo, P. et al. FLI1 is a novel ETS transcription factor involved in gene fusions in prostate cancer. Genes Chromosomes Cancer 51, 240–9 (2012).

6. Perner, S. et al. TMPRSS2-ERG fusion prostate cancer: an early molecular event associated with invasion. Am J Surg Pathol 31, 882–8 (2007).

7. Roudier, M.P. et al. Characterizing the molecular features of ERG-positive tumors in primary and castration resistant prostate cancer. The Prostate 76, 810–822 (2016).

8. Donaldson, L.W., Petersen, J.M., Graves, B.J. & McIntosh, L.P. Solution structure of the ETS domain from murine Ets-1: a winged helix-turn-helix DNA binding motif. The EMBO Journal 15, 125–134 (1996).

9. Wei, G.-H. et al. Genome-wide analysis of ETS-family DNA-binding in vitro and in vivo. The EMBO Journal 29, 2147–2160 (2010).

10. Wu, L. et al. ERG is a critical regulator of Wnt/LEF1 signaling in prostate cancer. Cancer Res 73, 6068–79 (2013).

11. Kron, K.J. et al. TMPRSS2-ERG fusion co-opts master transcription factors and activates NOTCH signaling in primary prostate cancer. Nat Genet 49, 1336–1345 (2017).

12. Yu, J. et al. An integrated network of androgen receptor, polycomb, and TMPRSS2-ERG gene fusions in prostate cancer progression. Cancer Cell 17, 443–54 (2010).

13. Sun, C. et al. TMPRSS2-ERG fusion, a common genomic alteration in prostate cancer activates C-MYC and abrogates prostate epithelial differentiation. Oncogene 27, 5348–5353 (2008).

14. Hollenhorst, P.C., Shah, A.A., Hopkins, C. & Graves, B.J. Genome-wide analyses reveal properties of redundant and specific promoter occupancy within the ETS gene family. Genes Dev 21, 1882–94 (2007).

15. Sandoval, G.J. et al. Binding of TMPRSS2-ERG to BAF Chromatin Remodeling Complexes Mediates Prostate Oncogenesis. Mol Cell 71, 554–566.e7 (2018).

16. Clark, J. et al. Diversity of TMPRSS2-ERG fusion transcripts in the human prostate. Oncogene 26, 2667–2673 (2007).

17. Gan, W. et al. SPOP Promotes Ubiquitination and Degradation of the ERG Oncoprotein to Suppress Prostate Cancer Progression. Mol Cell 59, 917–30 (2015).

18. An, J. et al. Truncated ERG Oncoproteins from TMPRSS2-ERG Fusions Are Resistant to SPOP-Mediated Proteasome Degradation. Mol Cell 59, 904–16 (2015).

19. Bose, R. et al. ERF mutations reveal a balance of ETS factors controlling prostate oncogenesis. Nature 546, 671–675 (2017).

20. Arora, V.K. et al. Glucocorticoid receptor confers resistance to antiandrogens by bypassing androgen receptor blockade. Cell 155, 1309–22 (2013).

21. Puhr, M. et al. The Glucocorticoid Receptor Is a Key Player for Prostate Cancer Cell Survival and a Target for Improved Antiandrogen Therapy. Clinical Cancer Research 24, 927–938 (2018).

22. Isikbay, M. et al. Glucocorticoid receptor activity contributes to resistance to androgen-targeted therapy in prostate cancer. Horm Cancer 5, 72–89 (2014).

23. Shah, N. et al. Regulation of the glucocorticoid receptor via a BET-dependent enhancer drives antiandrogen resistance in prostate cancer. eLife 6, e27861 (2017).

24. Li, J. et al. Aberrant corticosteroid metabolism in tumor cells enables GR takeover in enzalutamide resistant prostate cancer. Elife 6(2017).

25. Srivastava, S. et al. ETS Proteins Bind with Glucocorticoid Receptors: Relevance for Treatment of Ewing Sarcoma. Cell Rep 29, 104–117.e4 (2019).

26. Wilkinson, S. et al. Nascent Prostate Cancer Heterogeneity Drives Evolution and Resistance to Intense Hormonal Therapy. Eur Urol 80, 746–757 (2021).

27. Karzai, F. et al. Sequential Prostate Magnetic Resonance Imaging in Newly Diagnosed High-risk Prostate Cancer Treated with Neoadjuvant Enzalutamide is Predictive of Therapeutic Response. Clin Cancer Res 27, 429–437 (2021).

28. Montgomery, B. et al. Impact of baseline corticosteroids on survival and steroid androgens in metastatic castration-resistant prostate cancer: exploratory analysis from COU-AA-301. Eur Urol 67, 866–73 (2015).

29. Zhao, J.L. et al. The Effect of Corticosteroids on Prostate Cancer Outcome Following Treatment with Enzalutamide: A Multivariate Analysis of the Phase III AFFIRM Trial. Clin Cancer Res 28, 860–869 (2022).

30. Hussain, M. et al. Targeting Androgen Receptor and DNA Repair in Metastatic Castration-Resistant Prostate Cancer: Results From NCI 9012. J Clin Oncol 36, 991–999 (2018).

31. Sowalsky, A.G. et al. Neoadjuvant-Intensive Androgen Deprivation Therapy Selects for Prostate Tumor Foci with Diverse Subclonal Oncogenic Alterations. Cancer Res 78, 4716–4730 (2018).

32. Quigley, D.A. et al. Genomic Hallmarks and Structural Variation in Metastatic Prostate Cancer. Cell 175, 889 (2018).

33. Lundberg, A. et al. The Genomic and Epigenomic Landscape of Double-Negative Metastatic Prostate Cancer. Cancer Res 83, 2763–2774 (2023).

34. Xie, N. et al. The expression of glucocorticoid receptor is negatively regulated by active androgen receptor signaling in prostate tumors. Int J Cancer 136, E27–38 (2015).

35. Puhr, M. et al. The Glucocorticoid Receptor Is a Key Player for Prostate Cancer Cell Survival and a Target for Improved Antiandrogen Therapy. Clin Cancer Res 24, 927–938 (2018).

36. Michnick, S.W., Ear, P.H., Manderson, E.N., Remy, I. & Stefan, E. Universal strategies in research and drug discovery based on protein-fragment complementation assays. Nature reviews Drug discovery 6, 569–82 (2007).

37. Oakley, R.H. & Cidlowski, J.A. Homologous down regulation of the glucocorticoid receptor: the molecular machinery. Crit Rev Eukaryot Gene Expr 3, 63–88 (1993).

38. Tomlins, S.A. et al. Role of the TMPRSS2-ERG gene fusion in prostate cancer. Neoplasia 10, 177–88 (2008).

39. Hollenhorst, P.C., Paul, L., Ferris, M.W. & Graves, B.J. The ETS gene ETV4 is required for anchorage-independent growth and a cell proliferation gene expression program in PC3 prostate cells. Genes Cancer 1, 1044–1052 (2011).

40. Pellecchia, A. et al. Overexpression of ETV4 is oncogenic in prostate cells through promotion of both cell proliferation and epithelial to mesenchymal transition. Oncogenesis 1, e20 (2012).

41. Chai, X. et al. Computationally guided discovery of novel non-steroidal AR-GR dual antagonists demonstrating potency against antiandrogen resistance. Acta Pharmacol Sin 44, 1500–1518 (2023).

42. Hollenhorst, P.C., McIntosh, L.P. & Graves, B.J. Genomic and biochemical insights into the specificity of ETS transcription factors. Annual review of biochemistry 80, 437–71 (2011).

43. Frank, F., Ortlund, E.A. & Liu, X. Structural insights into glucocorticoid receptor function. Biochem Soc Trans 49, 2333–2343 (2021).

44. Hong, Z. et al. DNA Damage Promotes TMPRSS2-ERG Oncoprotein Destruction and Prostate Cancer Suppression via Signaling Converged by GSK3β and WEE1. Mol Cell 79, 1008–1023.e4 (2020).

45. Kirschke, E., Goswami, D., Southworth, D., Griffin, P.R. & Agard, D.A. Glucocorticoid receptor function regulated by coordinated action of the Hsp90 and Hsp70 chaperone cycles. Cell 157, 1685–97 (2014).

46. Centenera, M.M. et al. Co-targeting AR and HSP90 suppresses prostate cancer cell growth and prevents resistance mechanisms. Endocr Relat Cancer 22, 805–18 (2015).

47. Germain, L. et al. Preclinical models of prostate cancer -modelling androgen dependency and castration resistance in vitro, ex vivo and in vivo. Nat Rev Urol 20, 480–493 (2023).

48. Gerber, A.N., Newton, R. & Sasse, S.K. Repression of transcription by the glucocorticoid receptor: A parsimonious model for the genomics era. J Biol Chem 296, 100687 (2021).

49. Nhili, R. et al. Targeting the DNA-binding activity of the human ERG transcription factor using new heterocyclic dithiophene diamidines. Nucleic Acids Res 41, 125–38 (2013).

50. Regan, M.C. et al. Structural and dynamic studies of the transcription factor ERG reveal DNA binding is allosterically autoinhibited. Proc Natl Acad Sci U S A 110, 13374–9 (2013).

51. Lorenzin, F. & Demichelis, F. Past, Current, and Future Strategies to Target ERG Fusion-Positive Prostate Cancer. Cancers (Basel) 14(2022).

52. Khosh Kish, E., et al. The Expression of Proto-Oncogene ETS-Related Gene (ERG) Plays a Central Role in the Oncogenic Mechanism Involved in the Development and Progression of Prostate Cancer. Int J Mol Sci 23(2022).

53. Liberzon, A. et al. The Molecular Signatures Database (MSigDB) hallmark gene set collection. Cell Syst 1, 417–425 (2015).

54. Xiao, L. et al. Targeting SWI/SNF ATPases in enhancer-addicted prostate cancer. Nature 601, 434–439 (2022).

55. Fleseriu, M. & Petersenn, S. Medical therapy for Cushing’s disease: adrenal steroidogenesis inhibitors and glucocorticoid receptor blockers. Pituitary 18, 245–52 (2015).

56. Navone, N.M. et al. Movember GAP1 PDX project: An international collection of serially transplantable prostate cancer patient-derived xenograft (PDX) models. Prostate 78, 1262–1282 (2018).

57. Narayanan, S., Srinivas, S. & Feldman, D. Androgen-glucocorticoid interactions in the era of novel prostate cancer therapy. Nat Rev Urol 13, 47–60 (2016).

58. Srivastava, S. et al. ETS Proteins Bind with Glucocorticoid Receptors: Relevance for Treatment of Ewing Sarcoma. Cell Rep 29, 104–117 e4 (2019).

59. Yang, Z. et al. Corticosteroid switch from prednisone to dexamethasone in metastatic castration-resistant prostate cancer patients with biochemical progression on abiraterone acetate plus prednisone. BMC Cancer 21, 919 (2021).

60. Teo, M.Y. & Scher, H.I. Lessons from the SWITCH trial: changing glucocorticoids in the management of metastatic castration-resistant prostate cancer (mCRPC). Br J Cancer 119, 1041–1043 (2018).

61. Romero-Laorden, N. et al. Phase II pilot study of the prednisone to dexamethasone switch in metastatic castration-resistant prostate cancer (mCRPC) patients with limited progression on abiraterone plus prednisone (SWITCH study). Br J Cancer 119, 1052–1059 (2018).

62. Ndibe, C., Wang, C.G. & Sonpavde, G. Corticosteroids in the management of prostate cancer: a critical review. Curr Treat Options Oncol 16, 6 (2015).

63. Teply, B.A., Luber, B., Denmeade, S.R. & Antonarakis, E.S. The influence of prednisone on the efficacy of docetaxel in men with metastatic castration-resistant prostate cancer. Prostate Cancer Prostatic Dis 19, 72–8 (2016).

64. Bakour, N., Moriarty, F., Moore, G., Robson, T. & Annett, S.L. Prognostic Significance of Glucocorticoid Receptor Expression in Cancer: A Systematic Review and Meta-Analysis. Cancers (Basel) 13(2021).

65. Abida, W. et al. Phase I Study of ORIC-101, a Glucocorticoid Receptor Antagonist, in Combination with Enzalutamide in Patients with Metastatic Castration-resistant Prostate Cancer Progressing on Enzalutamide. Clin Cancer Res 30, 1111–1120 (2024).

66. Dobin, A. et al. STAR: ultrafast universal RNA-seq aligner. Bioinformatics 29, 15–21 (2013).

67. Love, M.I., Huber, W. & Anders, S. Moderated estimation of fold change and dispersion for RNA-seq data with DESeq2. Genome Biol 15, 550 (2014).

68. Benjamini, Y. & Hochberg, Y. Controlling the false discovery rate: a practical and powerful approach to multiple testing. Journal of the Royal Society Series B 57, 289–300 (1995).

69. Chen, E.Y. et al. Enrichr: interactive and collaborative HTML5 gene list enrichment analysis tool. BMC Bioinformatics 14, 128 (2013).

70. Blecher-Gonen, R. et al. High-throughput chromatin immunoprecipitation for genome-wide mapping of in vivo protein-DNA interactions and epigenomic states. Nat Protoc 8, 539–554 (2013).

71. Nguyen, H.M. et al. LuCaP Prostate Cancer Patient-Derived Xenografts Reflect the Molecular Heterogeneity of Advanced Disease an--d Serve as Models for Evaluating Cancer Therapeutics. Prostate 77, 654–671 (2017).

