## Supplemental figure and table for "*TMPRSS2-ERG* confers resistance to antiandrogens: mechanism and therapeutic implications"

**Supplementary figures, titles and legends (and Table S1)**

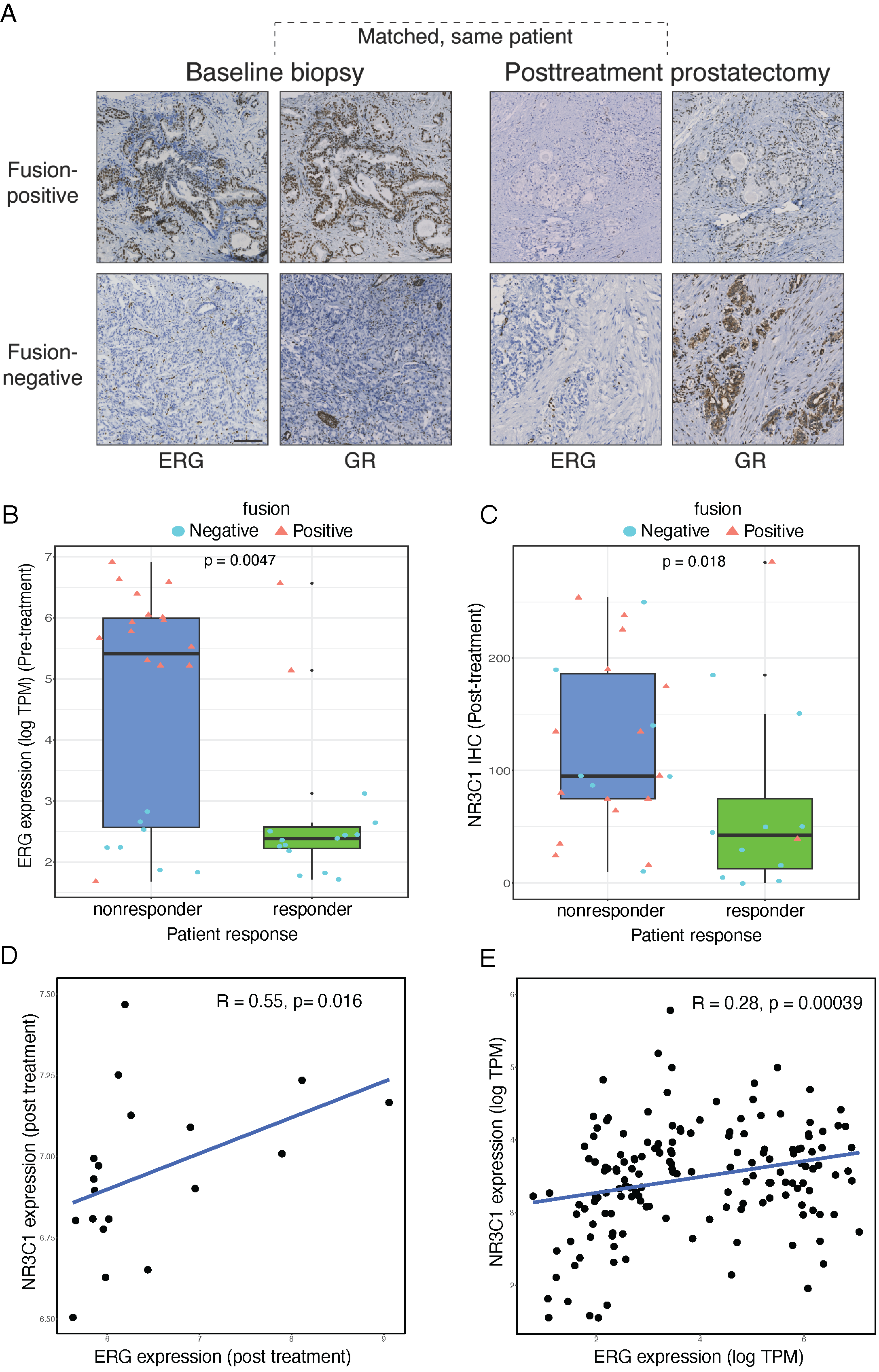

**Figure S1:  Pre- and post treatment immunohistochemical and statistical analyses indicate that tERG positivity and high GR (post-treatment) can predict resistance to ADT plus enzalutamide (related to Figure 1).** (**A**) Targeted biopsies were obtained from 37 men with intermediate- to high-risk prostate cancer before receiving ADT plus enzalutamide for 6 months. The tissues were used for IHC. Shown are representative micrographs of immunohistochemical (IHC) staining with anti-ERG and anti-GR antibodies that were applied on serial sections of matched baseline (left) and posttreatment (right) prostate tumor tissues from a TMPRSS2-ERG fusion-positive (top) and from a fusion-negative (bottom) case. Bar, 100 µm. (**B**) The patients from A were stratified according to the status of their pre-treatment ERG, either wild type or tERG. The boxplot presents pre-treatment ERG expression (in log TPM) vs. patient response. Fusion status of ERG in patients is shown as red for positive fusion (tERG) and cyan for negative fusion (wild type). Note that patients were grouped into non-responders or responders to ADT plus enzalutamide. P-values are from one-sided Wilcoxon tests. (**C**) Boxplot of post-treatment NR3C1 vs. patient responses. NR3C1 (GR) levels were determined using IHC post-treatment with ADT plus enzalutamide. Patients were grouped into responders and non-responders to the drug combination and the results were analyzed as in B. P-values are from one-sided Wilcoxon tests. (**C**) Spearman rank correlation between ERG and NR3C1 expression in 19 patients (from GSE102124) post treatment with abiraterone plus ADT. (**D**) Spearman rank correlation between ERG and GR (NR3C1) expression post-ADT treatment, based on analysis of RNA-seq data from 160 patients patients with PCa (WCDT dataset).

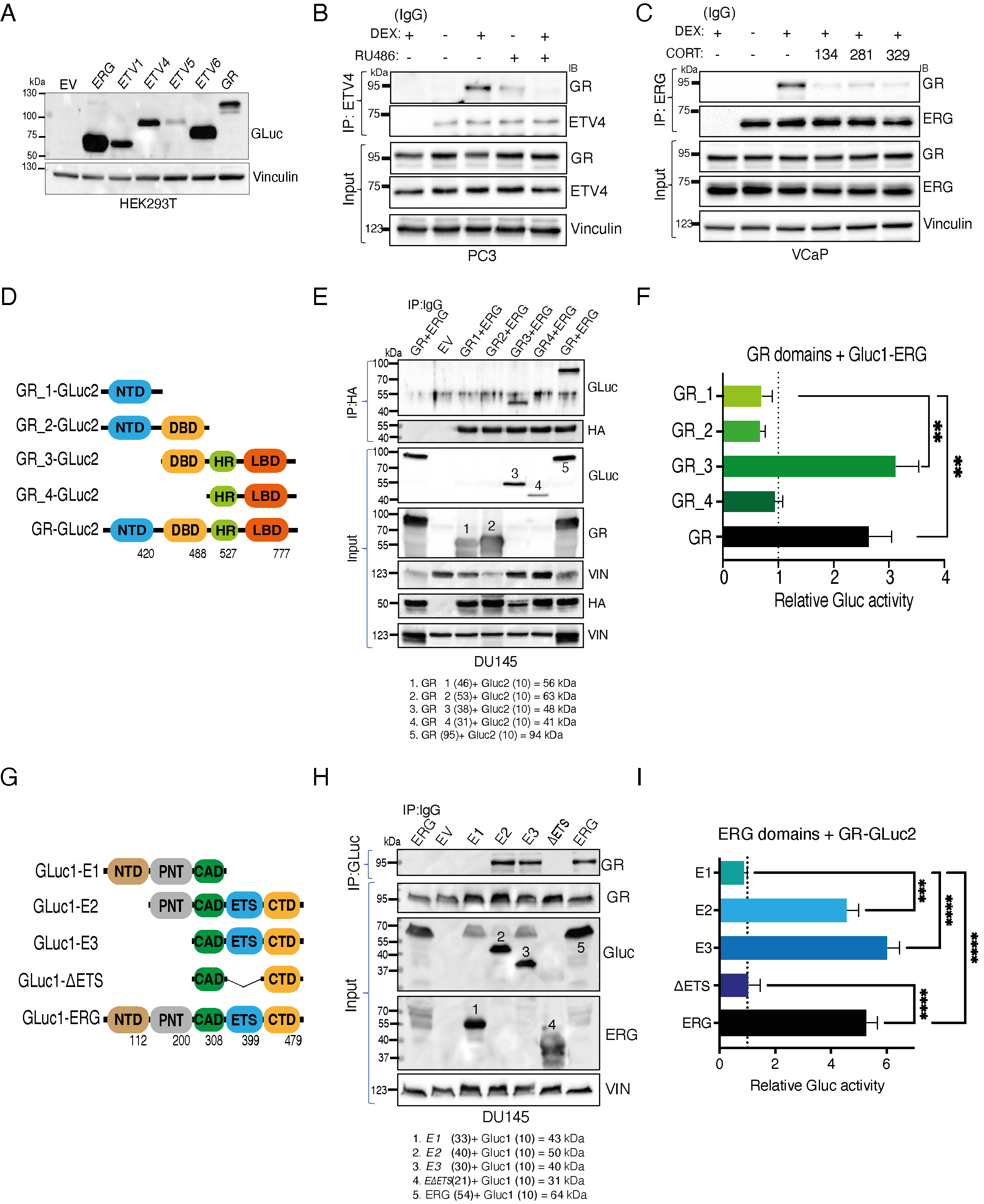

**Figure S2: The DBD-HR-LBD domain of GR interacts with the ETS domain of ERG (related to Fig. 2).** (**A**) An immunoblot showing the levels of the Gluc fusion proteins we transiently expressed in HEK293T cells (5x10^5^). Cells were harvested 24 hours post-transfection and processed for immunoblotting. Vinculin was used as the gel loading control. (**B**) Serum-starved PC3 cells (naturally overexpressing ETV4) were treated for 60 minutes with vehicle, DEX (1 μM), RU486 (1 μM), or the combination of drugs. Extracts were processed for co-immunoprecipitation (IP) and immunoblotting (IB). IgG, control antibody. (**C**) Serum-starved VCaP cells were treated for 60 minutes with vehicle (CON) or DEX (1 μM), in the absence or presence of the indicated SGRM compounds (C134, C281 and C329; 10 μM each). Whole cell extracts were processed for co-immunoprecipitation (IP) of ERG and GR, followed by immunoblotting (IB). (**D** and **E**) A schematic representation of the various domains of GR fused to Gluc2, along with their levels of expression in HEK293T cells. The indicated domains were inserted N-terminally to Gluc2. NTD, N-terminal domain; DBD, DNA binding domain; HR, hinge region; LBD, ligand binding domain. DU145 cells were transfected with Gluc1-ERG plasmids encoding different deletion mutants of ERG (see molecular weight calculations at the bottom of the panel). The Gluc1-ERG proteins were immunoprecipitated (IP) using a specific antibody. Immunoblotting (IB) was performed using antibodies that detected the endogenous forms of GR. (**F**) HEK293T cells (6x10^3^) were co-transfected with Gluc2 plasmids encoding different domains of GR and a Gluc1 plasmid encoding ERG (full length). Following 24 hours of incubation, cells were starved overnight and then treated for 60 minutes with vehicle, or with DEX (1 μM). The normalized luminescence activity of each construct is presented. (**G** and **H**) A schematic representation of the various domains of ERG fused to Gluc1, along with their levels of expression in transfected DU145 cells. The indicated domains were inserted C-terminally to Gluc1. NTD, N-terminal domain; PNT, pointed domain; CAD, central activation domain; ETS, DNA binding domain; and CTD, C-terminal domain. See cloning primer sequences in Table S1. (**I**) The Gluc2 plasmid encoding full length GR was co-transfected in DU145 cells together with the indicated Gluc1 plasmids encoding different domains of ERG. Following 24 hours of incubation, cells were starved overnight and then treated for 60 minutes with vehicle or with DEX (1 μM). Luminescence was determined in biological triplicates. All experiments were repeated three times. **, p< 0.01; ***, p< 0.001; ****, p < 0.0001. Note: the quantitative analyses (panels F and I) were repeated at least twice.

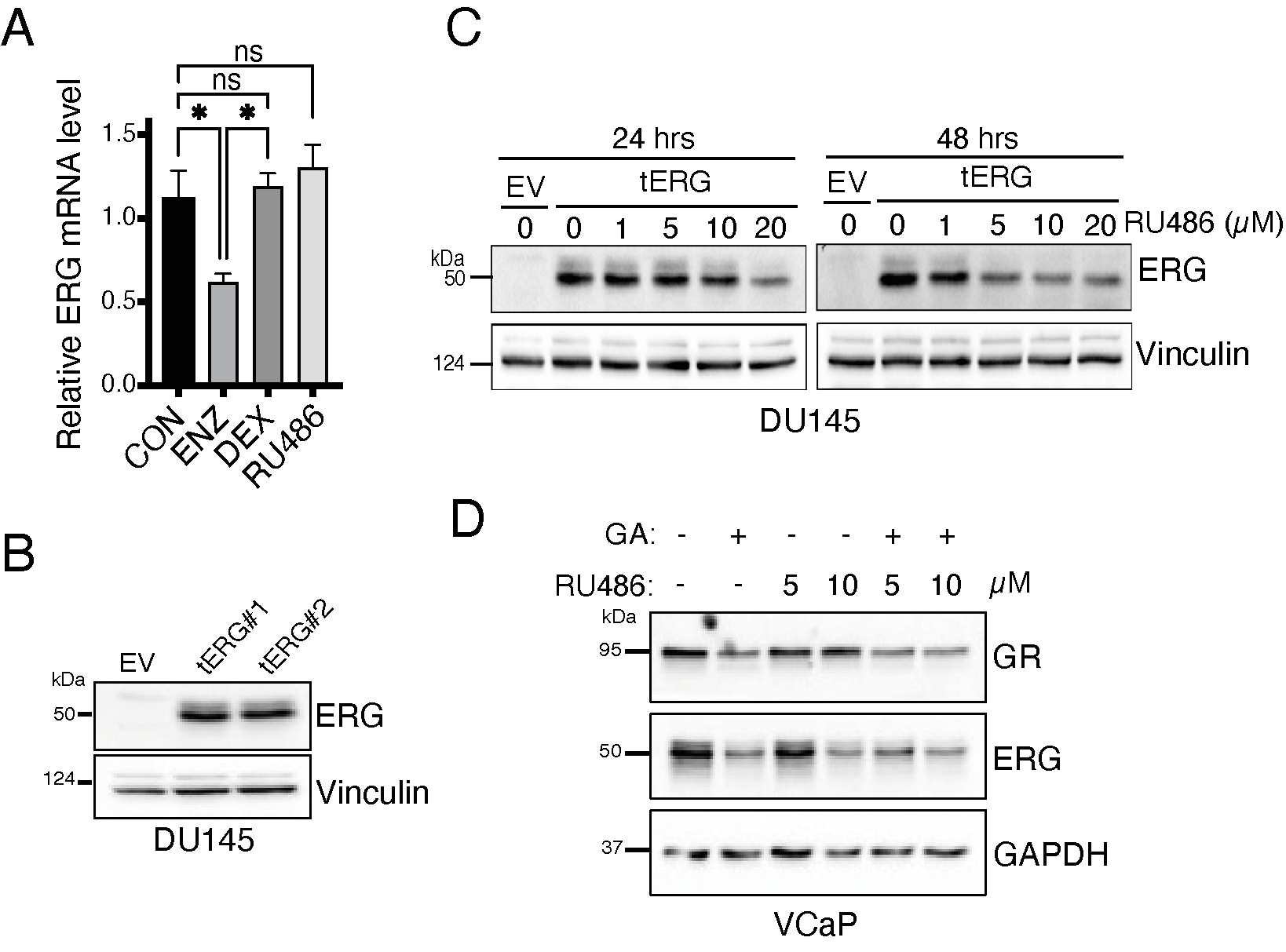

**Figure S3: Inhibition of GR destabilizes ERG (related to Fig. 3).** (**A**) VCaP cells were treated for 12, 24, 48 and 72 hours with the indicated concentrations of RU486 (either 5 or 10 μM) and, thereafter, whole cell extracts were probed for the endogenous ERG or vinculin. (**B**) DU145 cells were transfected with EV or tERG-encoding plasmids and selected with blasticidin for 10 days. The indicated two positive clones were used as the stable tERG over-expressing cells. (**C**) DU145 cells stably overexpressing tERG were treated for 24 or 48 hours with increasing concentrations of RU486. Thereafter, whole cell extracts were prepared and subjected to immunoblotting, as indicated. (**D**) VCaP cells were treated for 48 hours with geldanamycin (GA; 1μg/ml), RU486 (either 5 or 10 μg/ml) or with the combination of drugs. Whole cell extracts were prepared and subjected to immunoblotting using the indicated antibodies.

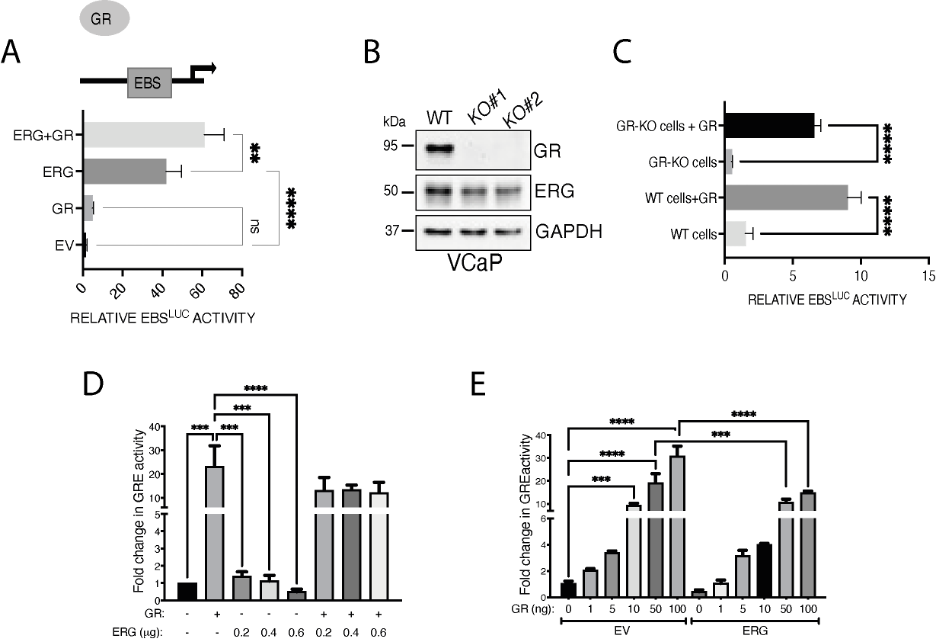

**Figure S4: Binding of GR with ERG boosts transcription from the EBS (ERG binding site; related to Fig. 4).** (**A**) HEK293T cells were co-transfected with an EBS-luciferase reporter plasmid, along with the indicated plasmids encoding ERG, GR, or the combination. Luciferase activity was determined 24 hours later using the Dual Luciferase Assay kit (from Promega). **, p< 0.01; ****, p < 0.0001; ns, not significant. (**B**) GR was stably knocked-out in VCaP cells using the CRISPR/Cas9 system and specific sgRNAs. Two cell clones were separately established. WT refers to cells that were transfected with a control guide RNA. Cell extracts were examined using immunoblotting for GR and ERG. GAPDH was used to ensure equal gel loading. (**C**) Wildtype VCaP cells or the respective GR-KO derivative cells were co-transfected with an EBS-luciferase plasmid. A GR-encoding plasmid, or a control vector, was co-transfected and 24 hours later we performed a luciferase assay, in triplicates. ****, p < 0.0001. (**D** and **E**) Serum-starved HEK293T cells were co-transfected with a GRE-luciferase promoter reporter, along with the indicated amounts of ERG and GR expression vectors. Twenty-four hours later, luciferase activity was measured using a luciferase assay kit.

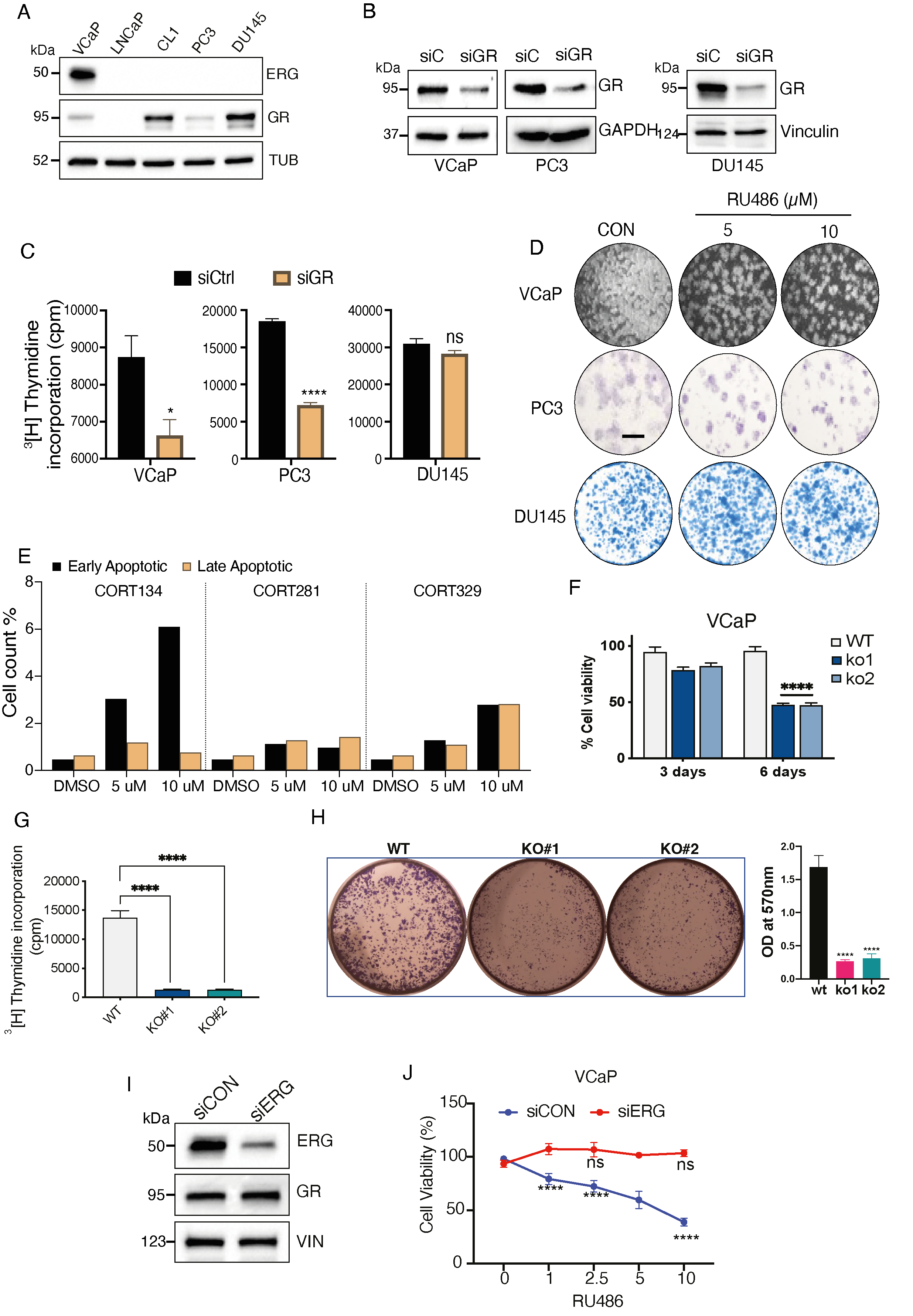

**Figure S5: Both genetic ablation and pharmacological inhibition of GR specifically associate with reduced proliferation of PCa cells expressing ETS fusions (related to Fig. 5).** (**A**) The indicated cell lines were cultured and then extracted. Whole extracts were subjected to immunoblotting for ERG and GR. Tubulin (TUB) was used to ensure equal gel loading. (**B** and **C**) VCaP, PC3 and DU145 cells were transfected with either control (scrambled) siRNAs (siCON) or GR-specific (siGR) oligonucleotides. Knockdown efficiency was monitored after 48 hours using immunoblotting with antibodies to GR. Cell proliferation was assayed by applying a radioactive thymidine incorporation assay. *, p < 0.05; ****, p < 0.0001; ns, not significant. (**D**) Shown are representative images corresponding to the colony formation assay and the bar plots shown in Figure 5C. Bar, 0.1 mm. (**E**) VCaP cells were seeded in 90-mm dishes. Thereafter, they were treated for 72 hours with the vehicle (DMSO) or with the indicated non-steroidal GR antagonists. Shown are results of an apoptosis assay performed in duplicates using an annexin V/7-AAD kit (from BioLegend). (**F**) WT and GR-knockout VCaP cells were seeded in 96-well plates and cell viability was measured using the XTT colorimetric assay following 3 or 6 days of incubation. (**G**) DNA synthesis was measured in WT and GR-KO VCaP cells (2 clones), using the radioactive thymidine incorporation assay. ****, p < 0.0001. (**H**) WT and GR-KO VCaP cells were sparsely seeded in 6-well plates. Fifteen days later, cells were fixed and stained with crystal violet. Photos are shown along with bar plots presenting the quantification of colonies. ****, p < 0.0001. (**I** and **J**) VCaP cells were transfected with either control oligonucleotides or with siRNAs targeting ERG. Forty-eight hours post transfection, the cells were harvested and subjected to immunoblotting (I). Alternatively, siCON- or siERG-transfected cells were seeded in 96-well plates and treated with the indicated concentrations of RU486, for 48 hours (J). Cell viability was measured using the XTT assay after 48 additional hours. ****, p < 0.0001; ns, not significant. Note: all quantitative analyses presented in this figure were repeated at least twice.

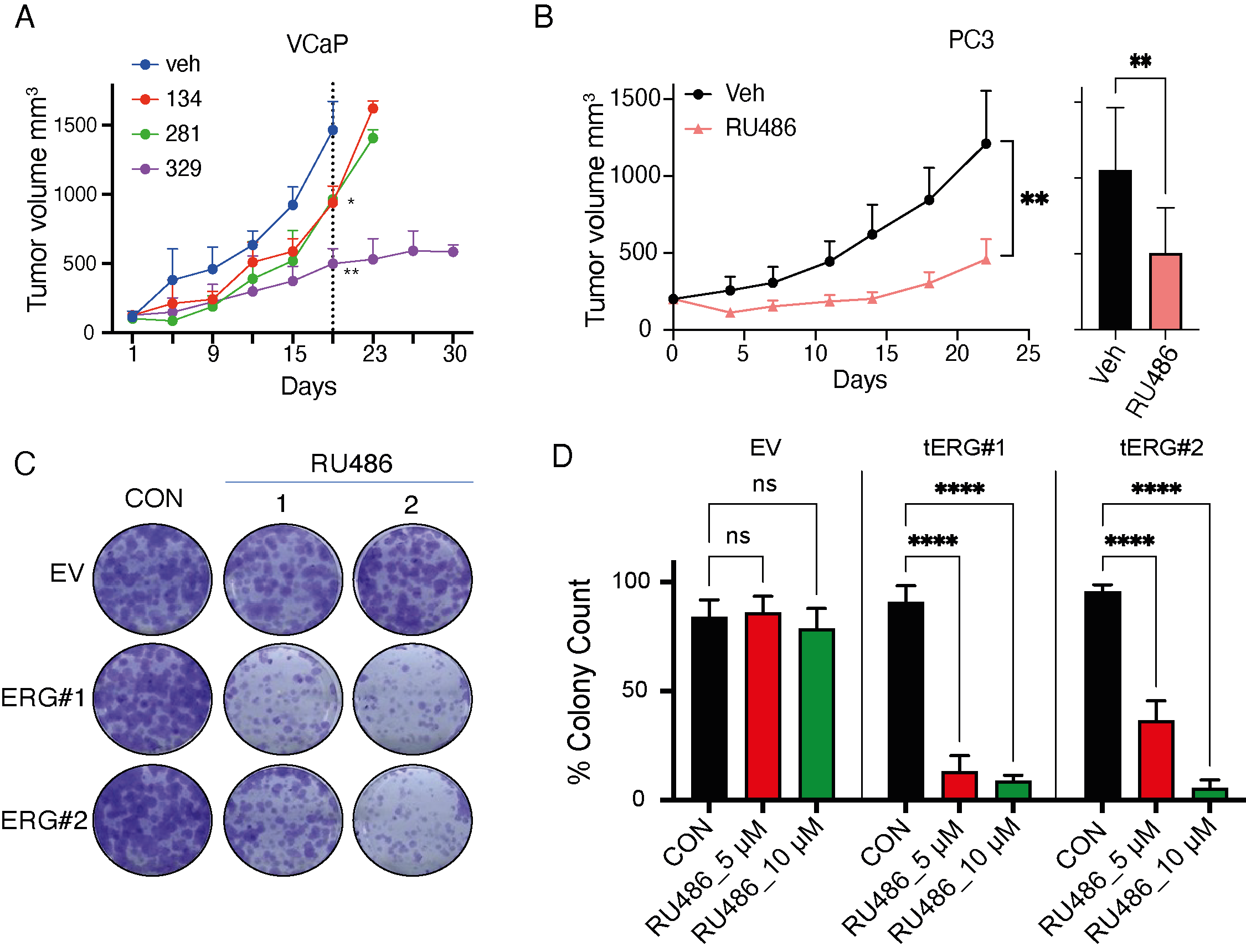

**Figure S6: GR inhibitors inhibit growth of ERG-expressing PCa cells but cells expressing low tERG do not respond to the inhibitors (related to Fig. 6).** (**A**) VCaP cells (2X10^6^) were implanted subcutaneously in athymic mice. Once tumors became palpable, animals were randomized into four groups (3 animals per group), which were daily treated with vehicle or with the indicated non-steroidal GR antagonists (50 mg/kg). The rates of tumor growth are shown. *, p < 0.05; **, p < 0.01. (**B**) PC3 cells (5X10^6^) were implanted in animals, which were randomized into groups that were daily treated with vehicle or with RU486 (1 mg/kg). The rates of tumor growth (left panel), along with tumor volumes on day 22 (bar plot), are shown. **, p < 0.01. (**C** and **D**) Two clones of DU145 cells stably expressing tERG were sparsely seeded in 6-well plates. Cells were later treated once every other day with either vehicle or RU486 (5 µM and 10 µM). Ten days later, all cells were fixed and stained with crystal violet. Representative photos are shown along with bar plots presenting the quantification of colony numbers in 5 non-overlapping microscope fields. The experiment was repeated twice. ****, p < 0.0001; ns, not significant.

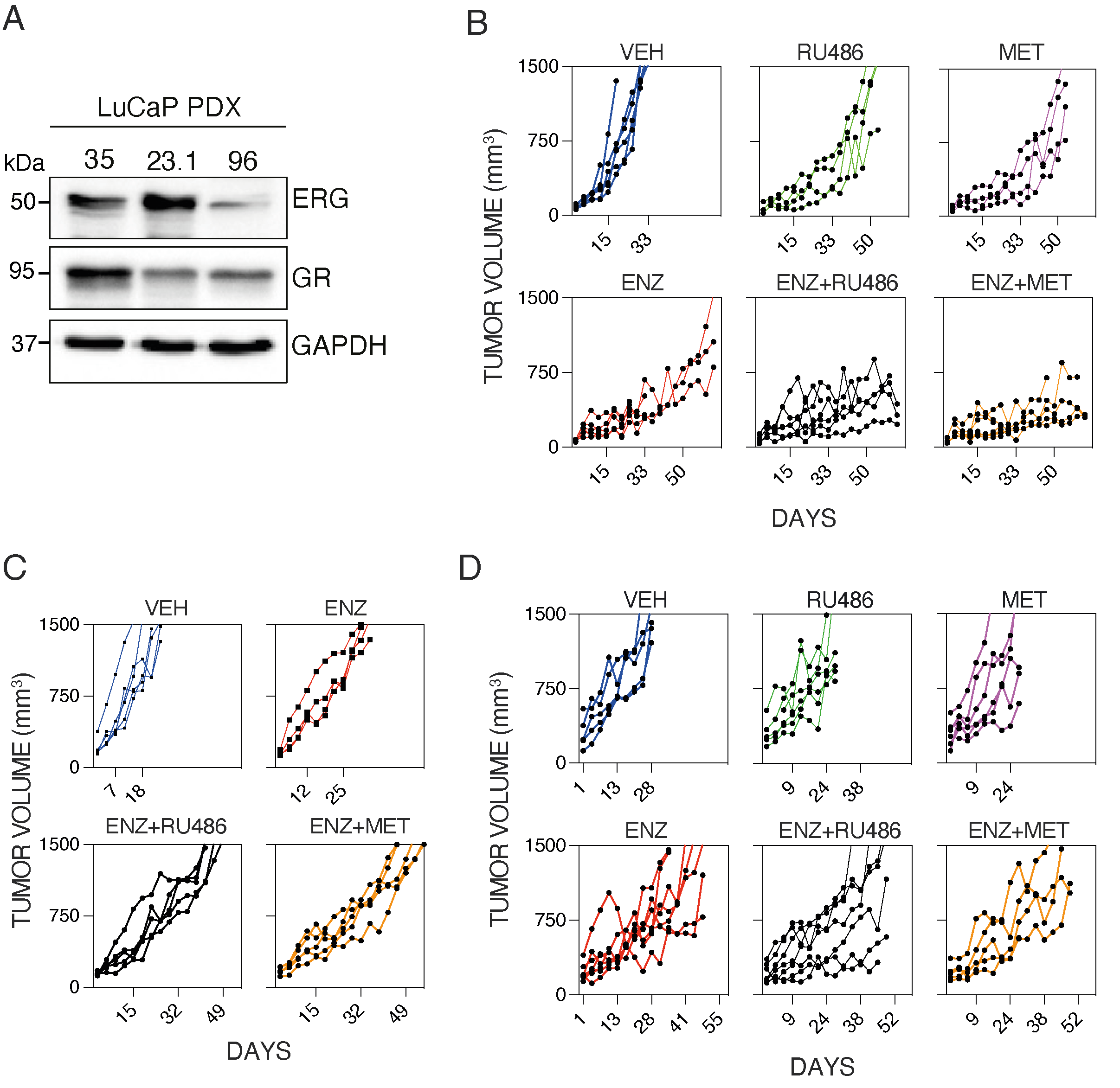

**Figure S7: Growth rates of three different PDX models in individual mice treated with combinations of GR and AR signaling inhibitors (related to Fig. 7).** (**A**) Whole extracts of the indicated PDX models were subjected to immunoblotting for ERG and GR. GAPDH was used to ensure equal gel loading. (**B**) Growth curves of individual tumors derived from the tERG-positive LuCaP 23.1 PDX model (see Fig. 7A). Each line corresponds to one animal. (**C**) Growth curves of individual tumors derived from the tERG-positive LuCaP 35 PDX model (see Fig. 7C). (**D**) Growth curves of individual tumors derived from the tERG-negative LuCaP 96 PDX model (see Fig. 7D).

| Cloning primers | Sequence |
| --- | --- |
| ERG-E1 forward | TGGTGGGTCCTCCGGAATTCAGACTGTCCCGGACCCA |
| ERG-E1 Reverse | AAACGGGCCCTCTAGATTAGGAGCTGTCCGACAGGAGCTC |
| ERG-E2 forward | TGGTGGGTCCTCCGGAAGCTACATGGAGGAGAAGCACATG |
| ERG-E2 Reverse | AAACGGGCCCTCTAGATTAGTAGTAAGTGCCCAGATGAGA |
| ERG-E3 forward | TGGTGGGTCCTCCGGACTCCACTACCTCAGAGAGACT |
| ERG-E3 Reverse | Same as E2 reverse |
| ΔETS forward | Same as E3 forward (used ΔETS plasmid as template ) |
| ΔETS Reverse | Same as E3 reverse (used ΔETS plasmid as template ) |

**Supplementary Table S1:** List of all primers used for DNA cloning and validation.
